# An atlas of the *C. elegans* starvation response reveals that transcription is necessary to initiate transcriptional silencing in quiescent primordial germ cells

**DOI:** 10.64898/2026.09.04.749554

**Authors:** Jingxian Chen, Sharan Surya, Rojin Chitrakar, Jenna R. Valley, Marcos Francisco Perez, L. Ryan Baugh

## Abstract

Starvation profoundly impacts development, but how different tissues respond to starvation is unclear. We generated a cellular atlas of gene expression in fed and starved *C. elegans* L1 larvae. We identified 79 cell types in both conditions, and we detected ∼98% of protein-coding genes, 44% of which were differentially expressed. Starvation affected translation-related genes across the animal, and most ‘housekeeping’ genes were downregulated. However, tissue-specific patterns of differential expression, transcription factor activity, and GO term enrichments are widespread, suggesting tissue-specific gene regulatory mechanisms and functional consequences. We inferred tissue-specific effects on transcription factor activity, identifying known and novel putative nutrient-dependent transcriptional regulators. Surprisingly, we found extensive transcription in starved primordial germ cells (PGCs), which are known to hyper-compact their chromatin and silence transcription during L1 starvation. We validated PGC transcription of *aak-1/AMPK*, and we showed that zygotic *aak-1/AMPK* is required for PGC chromatin hyper-compaction, supporting reproductive success upon recovery. This work provides a valuable resource characterizing nutritional control of transcription in an animal, and it reveals that transcription of *aak-1/AMPK* is necessary to silence transcription in quiescent PGCs.

## INTRODUCTION

Animals must sense and respond to fluctuations in nutrient availability to maintain developmental homeostasis. In well-fed animals, growth, cell proliferation, and reproduction are prioritized, but when nutrient availability is scarce, those processes are deprioritized in favor of survival. Some animals can enter a diapause-like state to endure starvation, which is characterized by arrested development and profound alterations to metabolism (Easwaran and Montell, 2023). When the nematode *Caenorhabditis elegans* hatches without food, they arrest postembryonic development in the first larval stage (L1 arrest or L1 diapause), and they can survive arrest for weeks (Baugh, 2013). During L1 arrest, cell proliferation, migration, fusion, and differentiation are halted (Baugh and Sternberg, 2006; Fukuyama *et al*, 2006), and conserved pathways that promote development are inhibited while those that support survival are activated (Baugh and Hu, 2020; Hibshman *et al*, 2017; Kaplan *et al*, 2015). *C. elegans* L1 arrest provides a powerful animal model to study nutritional control of gene regulation and development.

The anatomical complexity of the metazoan starvation response is unclear. A variety of nutrient-sensing mechanisms are conserved between metazoans and yeast (Chantranupong *et al*, 2015), suggesting that individual animal cells can respond autonomously to starvation. On the other hand, physiological constraints imposed by differentiation and specialized cellular function could cause different tissues to respond to starvation in different ways. Tissue-specific rescue of mutants affecting the FoxO transcription factor (TF) *daf-16*, the AMP-activated kinase AMPK, and the tumor suppressors *daf-18/PTEN* and *lin-35/Rb* suggests that these genes function in specific tissues and organs to promote developmental arrest and starvation resistance (Cui *et al*, 2013; Fukuyama *et al*, 2012; Kaplan *et al*, 2015). Determining the extent to which the starvation response is universal vs. tissue-specific requires systematic investigation of the response with cellular resolution.

The magnitude and dynamics of the *C. elegans* starvation response have been characterized, and regulatory mechanisms have been identified (Baugh and Hu, 2020). Most detected genes are differentially expressed in fed vs. starved L1 larvae (Baugh *et al*, 2009; Fisher *et al*, 2026; Webster *et al*, 2018), with the starvation response peaking within 12 h (Webster *et al*, 2022). During L1 arrest, RNA Polymerase II (RNAPII) is poised at the promoters of genes that support growth (Maxwell *et al*, 2014), and the starvation response is largely reversed within 1 h of feeding (Maxwell *et al*, 2012). Alternative mRNA splicing is regulated by nutrient availability (Maxwell *et al*, 2012), as is translation, with rapid up-regulation of translation of ribosomal proteins upon feeding (Stadler and Fire, 2013). These findings were all made with whole-animal (bulk) genomic analysis, but imaging has revealed tissue-specific mechanisms. Chromatin is reorganized in intestinal nuclei during starvation, supporting expression of metabolic and stress-related genes (Al-Refaie *et al*, 2024). In contrast, chromatin is hyper-compacted in primordial germ cells (PGCs) during L1 arrest to support global transcriptional quiescence and reproductive fitness (Belew *et al*, 2021; Webster *et al*, 2022).

Several conserved signaling, gene regulatory, and metabolic pathways that govern L1 arrest have been identified (Baugh and Hu, 2020). Approximately 100 genes are reported to affect starvation survival in *C. elegans*, and most have analogous effects on lifespan in fed adults. However, the anatomical site(s) of action of relatively few of these genes has been investigated (Baugh and Hu, 2020). A variety of transcriptional regulators are known to mediate the starvation response and affect starvation survival. The FoxO transcription factor DAF-16 is the primary effector of insulin/IGF signaling (IIS) during L1 starvation and is required for reduction of IIS to extend adult lifespan (Fisher *et al*, 2026; Kenyon *et al*, 1993). *daf-16/FoxO* promotes L1 arrest via repression of pathways that drive development (Kaplan *et al*, 2015), and it supports starvation survival via regulation of central carbon metabolism (Hibshman *et al*, 2017). *hlh-30/TFEB* collaborates with *daf-16/FoxO* during starvation, though it affects expression of more genes than *daf-16/FoxO* and is more sensitive to starvation (Munoz-Barrera *et al*, 2025; O’Rourke and Ruvkun, 2013; Settembre *et al*, 2013). *hlh-30/TFEB* also supports adult longevity (Lapierre *et al*, 2013). The nuclear hormone receptor *nhr-49/HNF4* supports starvation survival via transcriptional regulation of lipid metabolism, and it supports adult longevity (Goh *et al*, 2018; Van Gilst *et al*, 2005). *daf-18/PTEN*, *lin-35/Rb*, and the DREAM complex repress germline gene expression (Fry *et al*, 2021; Petrella *et al*, 2011; Wu *et al*, 2012), supporting starvation survival (Chen *et al*, 2025; Cui *et al*, 2013; Fukuyama *et al*, 2012), and they each promote longevity (Knutson *et al*, 2016; Ogg and Ruvkun, 1998). *pha-4/FoxA* supports starvation survival (Wu *et al*, 2018; Zhong *et al*, 2010), and *skn-1/Nrf* may also affect starvation survival (Paek *et al*, 2012). Notably, *pha-4/FoxA* and *skn-1/Nrf* are both required for dietary restriction to increase adult lifespan (Bishop and Guarente, 2007; Panowski *et al*, 2007). The bZIP transcription factor-encoding genes *zip-2* and *cebp-2* along with *cep-1/p53* surveil mitochondrial function and regulate DNA repair supporting innate immunity, starvation survival, and adult longevity (Derry *et al*, 2001; Hahm *et al*, 2019; Ventura *et al*, 2009; Yan *et al*, 2025). Nonetheless, additional factors likely contribute to transcriptional regulation of the starvation response.

Here we present the first atlas of the metazoan starvation response based on single-cell RNA sequencing (scRNA-seq), focusing on fed and starved *C. elegans* L1 larvae. Genes related to ribosome biogenesis and the nucleolus are affected across most tissues, suggesting a pervasive effect on translation. However, most genes are expressed in more tissues than they are differentially expressed in, and many genes are differentially expressed in a single tissue, suggesting tissue-specific regulation. Although primordial germ cells (PGCs) undergo chromatin hyper-compaction and global transcriptional silencing during L1 arrest, we observed widespread transcriptional activation in PGCs of larvae that recently hatched without food. We validated transcription of the AMPK catalytic α-subunit *aak-1* in PGCs with single-molecule fluorescence *in situ* hybridization (smFISH) and showed that it is required zygotically for PGC chromatin hyper-compaction and reproductive success upon recovery. This work provides a valuable reference dataset for nutritional control of development and gene regulation in a powerful animal model, and it provides mechanistic insight into nutrient-dependent regulation of chromatin structure and gene expression in quiescent PGCs.

## RESULTS

### A cellular atlas of gene expression in fed and starved L1 larvae

We used scRNA-seq in fed and starved wild-type (N2) L1 larvae to profile gene expression. We collected nine and eight replicates each of larvae that had hatched in the presence or absence of food (*E. coli*) ∼6 or ∼12 h earlier (‘fed’ and ‘starved’, respectively; Fig. 1A and Sup. Fig. 1). We chose 6 h for fed since the response to food is maximal by then (Maxwell *et al*, 2012) but few if any postembryonic cell divisions have occurred (Sulston and Horvitz, 1977). We chose 12 h for starved since that is when the starvation response peaks (Baugh *et al*, 2009; Webster *et al*, 2022). After filtering out low-quality cells and multiplets (Sup. Fig. 2), we obtained 52,278 and 50,902 single cells for fed and starved conditions, respectively (Sup. Data 1). Replicate samples were very well correlated (Sup. Fig. 3A), and the first principal component clearly separates fed and starved samples (Sup. Fig. 3B), explaining 82% of the variance and showing that the fed samples did not experience appreciable starvation during dissociation. For fed and starved, respectively, we detected 27,714 and 27,761 genes, or ∼59% of the annotated transcriptome, and 19,588 and 19,596 protein-coding genes, or ∼98% of protein-coding genes. Detection of a higher proportion of protein-coding genes is likely due at least in part to them having polyadenylated tails since an oligo-(dT) primer was used to prepare cDNA.

**Figure 1.**
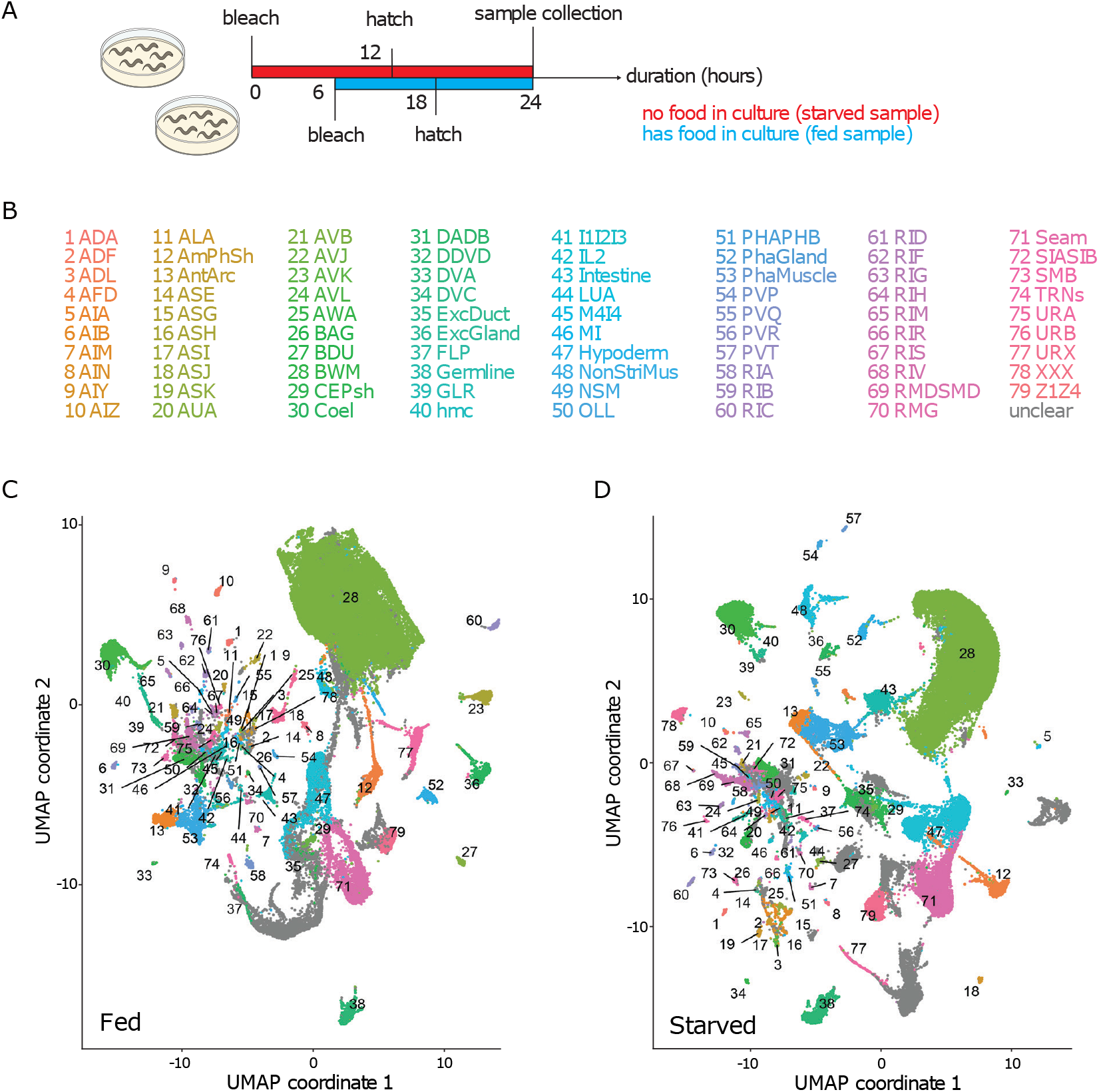
A cellular atlas of gene expression in fed and starved L1 larvae. (A) Schematic of experimental set-up. Starved is indicated with red and fed with blue. (B) A key for the 79 cell types shared between starved and fed conditions for C and D. Numbers and names have the same color as the cells (dots) they represent in C and D. (C, D) Fed and starved, respectively, UMAPs with cell types labelled by numbers in B. See also Sup. Fig. 1-3.

The uniform manifold approximation and projection (UMAP) algorithm (Becht *et al*, 2018) enabled visualization of clusters of cells in two dimensions (Fig. 1 B-D). We clustered fed and starved data independently. For fed cluster annotation, we used marker genes reported in previous scRNA-seq studies (Cao *et al*, 2017; Packer *et al*, 2019; Taylor *et al*, 2021) and other miscellaneous publications (Sup. Data 2). If there was no reported marker gene to confidently annotate a cluster, then we queried the most enriched genes in this cluster against previous scRNA-seq studies (Cao *et al*, 2017; Packer *et al*, 2019; Taylor *et al*, 2021). Only when the querying result unambiguously suggested a cluster identity did we use it for annotation. For starved cluster annotation, we followed the same method except that in addition to previous scRNA-seq data (Cao *et al*, 2017; Packer *et al*, 2019; Taylor *et al*, 2021), we used our fed scRNA-seq data for querying. We identified 93 and 104 cell types for fed and starved, respectively, with 79 cell types in common (Fig. 1B-D; Sup. Data 2).

### Differential expression in response to nutrient availability

We performed differential expression analysis between starved and fed conditions for each of the 79 shared cell types (‘upregulation’ implies higher expression in starved than fed hereafter). 8,777 total genes were differentially expressed (FDR < 0.05 and percentage of cells in the cluster with expression detected >= 10% in fed or starved), including 8,573 protein-coding genes (Sup. Data 3). Expression of individual genes can be queried with a web-based tool (https://jingxianchen.shinyapps.io/l1_sc/). Pseudobulk differential expression is concordant with published bulk analysis, validating our results (Sup. Fig. 4A). The non-coding RNA *tts-1* (<u>T</u>ranscribed <u>T</u>elomerase-like <u>S</u>equence), which represses translation to promote longevity (Essers *et al*, 2015), displayed the largest magnitude response, being upregulated in all cell types by an average of 97-fold. The remainder of our analysis will focus on protein-coding genes (hereafter ‘genes’). The median number of differentially expressed genes (DEGs) was 284 per cell type, with a range of 29 (AIY neurons) to 2,438 (intestine) (Fig. 2A). The low number of DEGs in AIY is not necessarily due to there being only two of these cells in the animal, since other cell types with only two cells had substantially more DEGs (*e.g.*, other neurons, somatic gonad precursors (Z1Z4), and germline (Z2Z3)). These results suggest that cells vary in their responsiveness to starvation.

**Figure 2.**
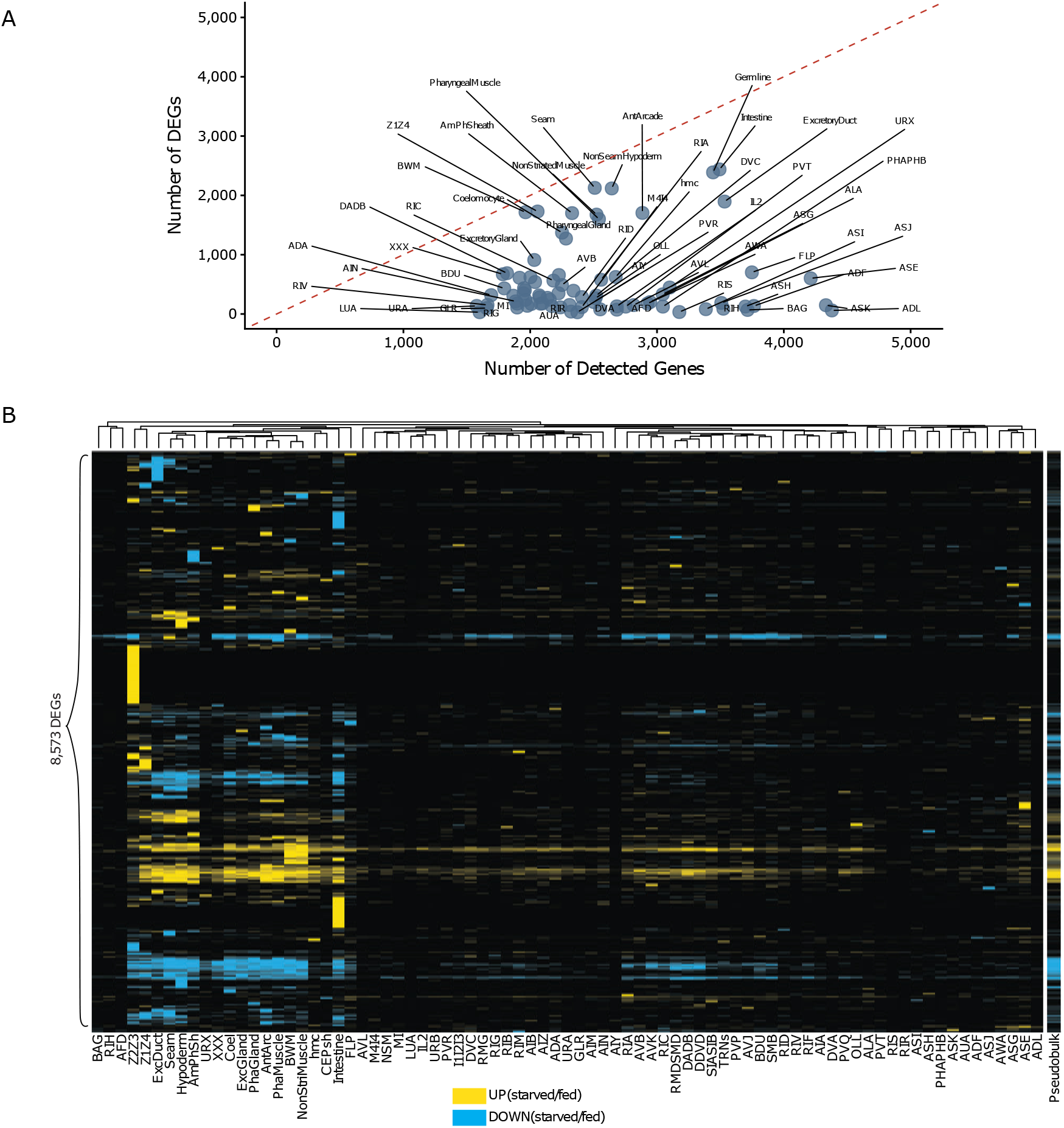
Cell type-specific differential expression in fed and starved L1 larvae. (A) Number of detected genes plotted against the number of DEGs for each of 79 cell types identified in both fed and starved larvae. The dashed red line indicates y = x. (B) Heatmap of genes and cell types showing up- and down-regulation of 8,573 DEGs differentially expressed in at least one cell type, as well as their differential expression in pseudobulk. Genes upregulated in starved compared to fed conditions are colored yellow, and downregulation is colored blue – degree of up or downregulation is not indicated. Heatmap rows and columns were hierarchically clustered and reordered by optimal leaf ordering (OLO). Only the cell-type dendrogram is shown. See also Sup. Fig. 4.

Few genes were differentially expressed in all cells they were detected in, with 3,422 (∼40%) DEGs differentially expressed in only one cell type and most differentially expressed in no more than two (Sup. Fig. 4B). We clustered DEGs and cell types to reveal anatomical patterns of regulation (Fig. 2B). There are prominent clusters of genes regulated exclusively in intestine or germline (Z2Z3), and their differential expression was not evident in pseudobulk. These observations suggest cell type-specific responses to starvation. In contrast, there are clusters of genes up or downregulated across many cell types, and their differential expression is evident in pseudobulk (Fig. 2B). Together these results suggest a combination of common and cell type-specific responses to starvation.

Functionally related cell types displayed less variation in their response to starvation. For example, germline (Z2Z3) and somatic gonad precursors (Z1Z4) clustered, as did non-striated muscle (NonStriMus) and body-wall muscle (BWM), as well as syncytial hypodermis (Hypoderm) and hypodermal seam cells (Seam) (Fig. 2B). Moreover, the chemosensory neurons AWA, ASG, ASE, and ADL, and the thermosensory neurons BAG, RIH, and AFD, each cluster together (Fig. 2B), suggesting related starvation responses among neurons of the same functional class.

We aggregated data from cell type to tissue level for less granular analysis. We defined eighteen tissues common to fed and starved (Fig. 3A-B, see Sup. Data 2 for aggregation of cell types into tissues). *C. elegans* L1 larvae have an invariant anatomy (Sulston and Horvitz, 1977), and there is a significant correlation between the number of expected and observed cells for each tissue in each condition, though some tissues are relatively over or under-represented (Sup. Fig. 5). We detected 7,646 DEGs in one or more tissues (FDR < 0.05 and percentage of cells in tissue with expression detected >= 10% in fed or starved) (Sup. Data 4). The median number of DEGs was 1,666 per tissue, with a range of 99 (GLR) to 2,438 (intestine) (Fig. 4A). ∼42% (3,215) of DEGs were differentially expressed in a single tissue (Fig. 4B), supporting the conclusion that much of the starvation response is cell/tissue-specific.

**Figure 3.**
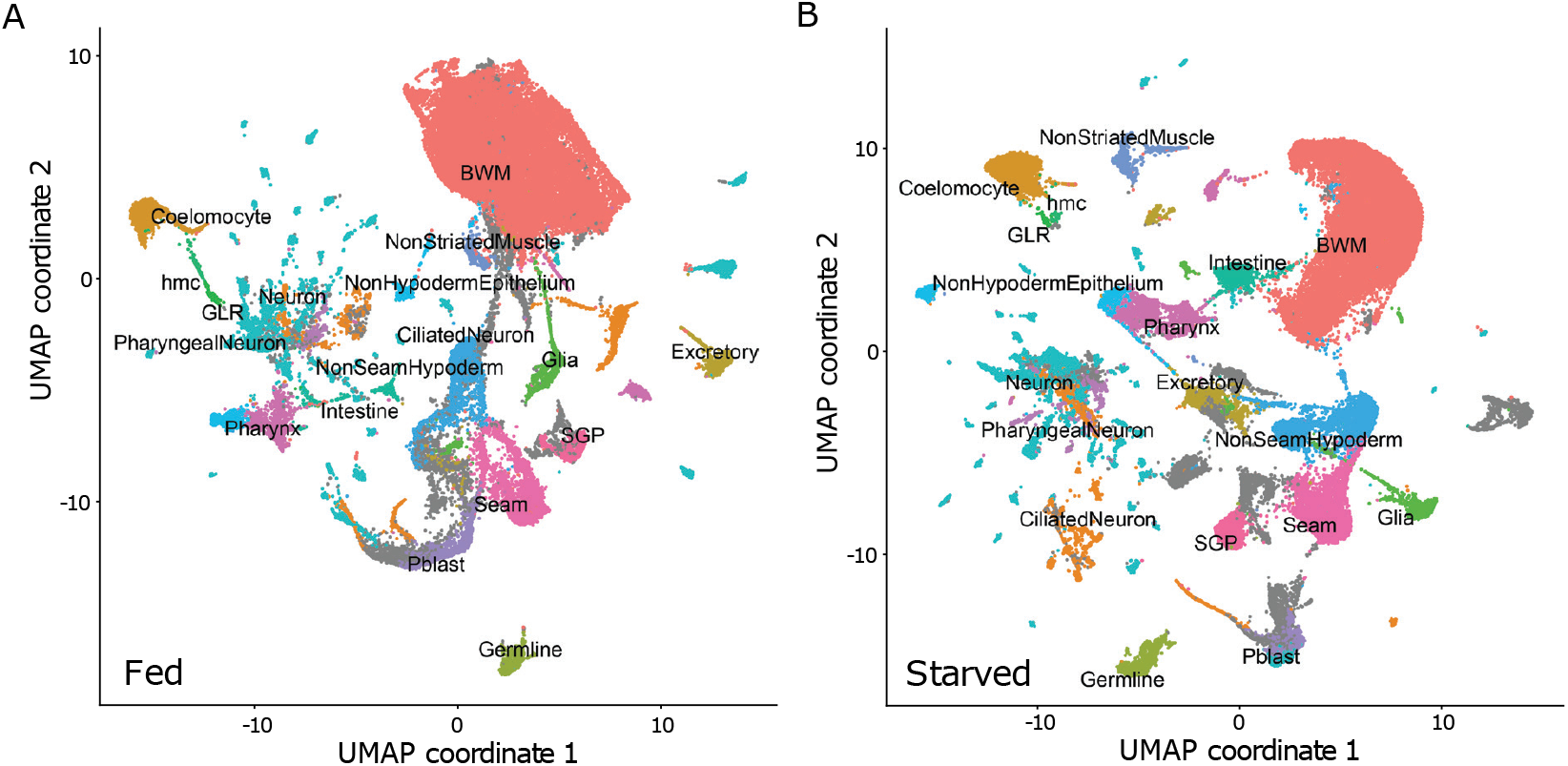
A tissue atlas of gene expression in fed and starved L1 larvae. (A, B) Fed and starved, respectively, UMAPs with tissue names labelled. Tissues have the same colors in both panels. See also Sup. Fig. 5.

**Figure 4.**
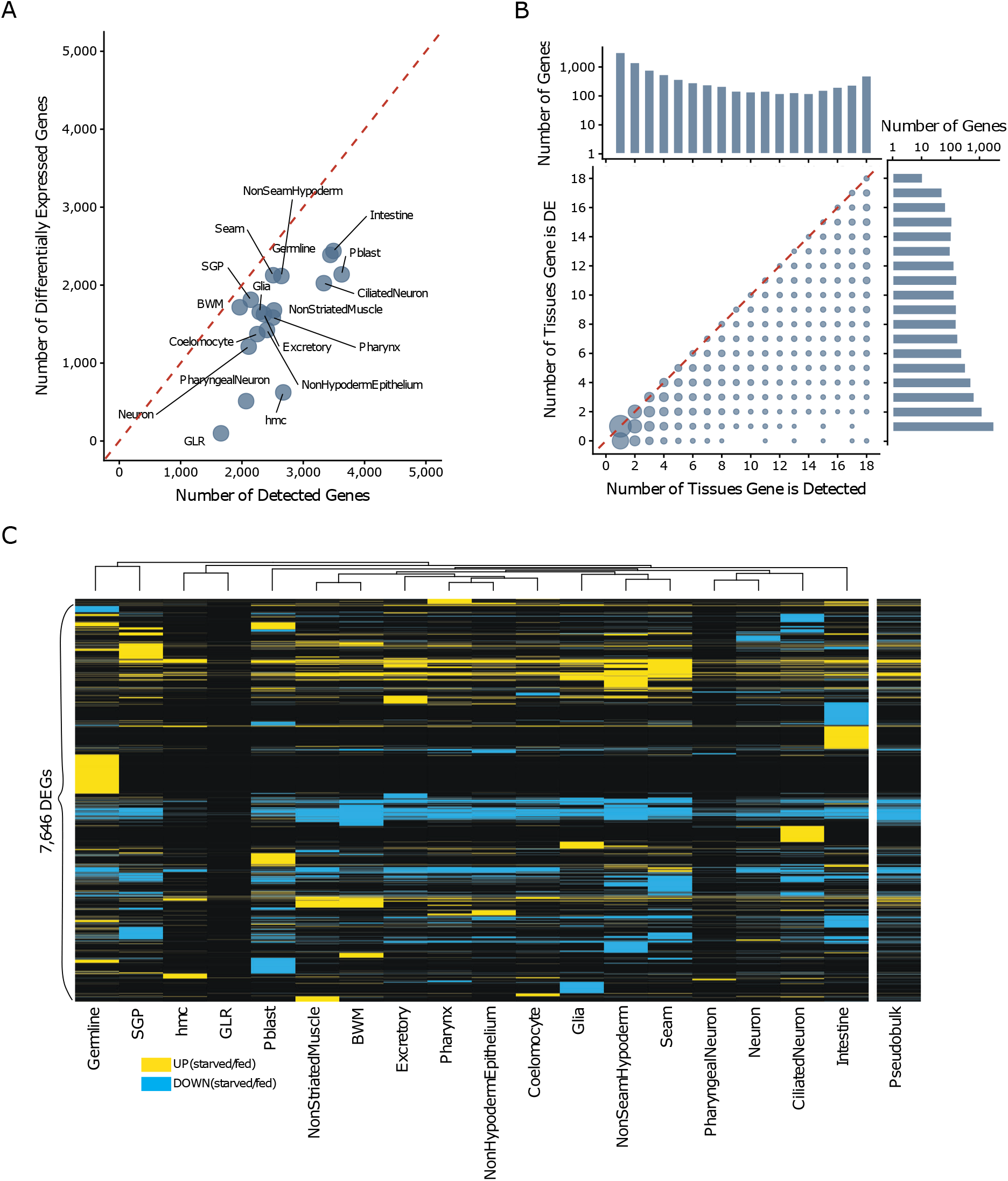
Tissue-specific differential expression in fed and starved L1 larvae. (A) Number of detected genes plotted against number of DEGs per tissue. (B) Bubble plot relating distributions of the number of tissues genes are detected in and the number of tissues genes are differentially expressed in. A one-dimensional histogram is plotted separately for each axis. (A, B) Dashed red line indicates y = x. (C) Heatmap showing up and downregulation of 7,646 DEGs differentially expressed in at least one tissue, as well as their differential expression in pseudobulk. Genes upregulated in starved compared to fed conditions are colored yellow, and downregulation is colored blue – degree of up or downregulation is not indicated. Heatmap rows and columns were hierarchically clustered, and rows were reordered by optimal leaf ordering (OLO). Only the tissue dendrogram is shown here. Column (tissue) order in panel C is used in figures hereafter. See also Sup. Fig. 6.

Clustering tissues and DEGs revealed tissue-specific starvation responses (Fig. 4C). In addition to the germline and intestine-specific clusters of DEGs seen in the cell-level analysis (Fig. 2B), ciliated neurons (CiliatedNeuron), ventral cord blast cells (Pblast), glia (Glia), and syncytial hypodermis (NonSeamHypoderm) have relatively many genes that are differentially expressed exclusively in that tissue (Sup. Fig. 6). Furthermore, many DEGs behaved differently in different tissues. Out of 4,431 genes differentially expressed in more than one tissue, only ∼60% (2,669) showed concordant effects across the affected tissues.

**Table 1.**
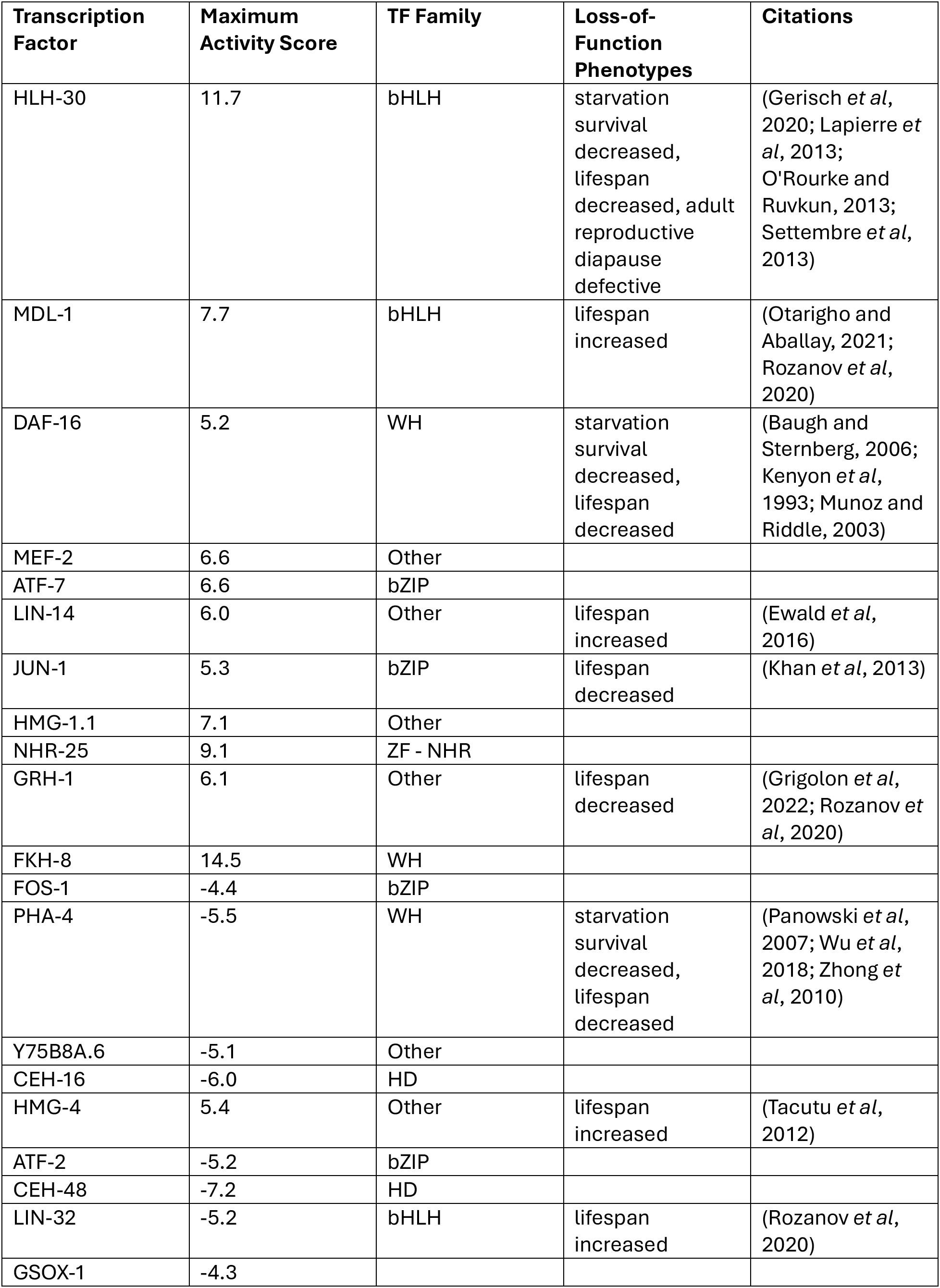
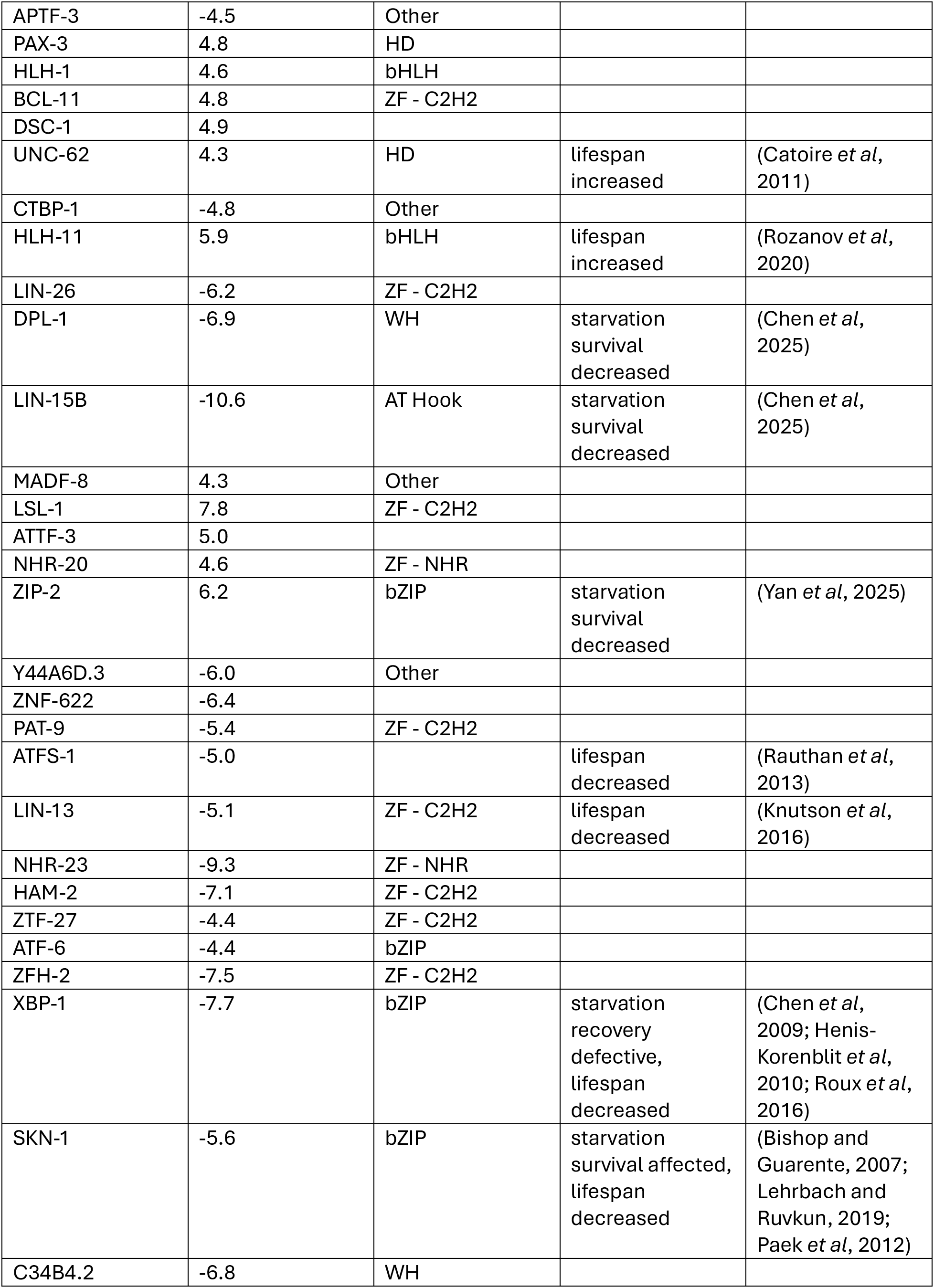
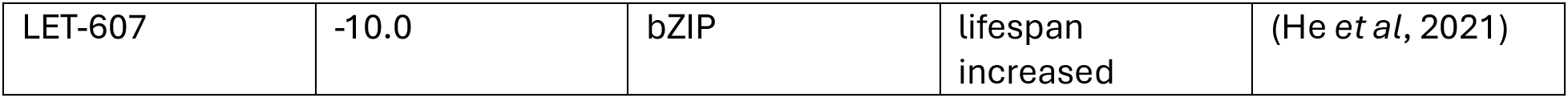
The top 50 TFs identified by *Cel*EsT as mediators of nutrient-dependent transcriptional regulation. TFs were selected based on the maximum absolute value of their tissue-specific activity scores. Activity scores, TF families, loss-of-function phenotypes, and supporting citations are provided. For phenotypes, only those related to starvation survival, fed adult lifespan, and adult reproductive diapause (starvation-induced developmental arrest in adults) are included.

| Transcription Factor | Maximum Activity Score | TF Family | Loss-of-Function Phenotypes | Citations |
| --- | --- | --- | --- | --- |
| HLH-30 | 11.7 | bHLH | starvation survival decreased, lifespan decreased, adult reproductive diapause defective | (Gerisch *et al*, 2020; Lapierre *et al*, 2013; O'Rourke and Ruvkun, 2013; Settembre *et al*, 2013) |
| MDL-1 | 7.7 | bHLH | lifespan increased | (Otarigho and Aballay, 2021; Rozanov *et al*, 2020) |
| DAF-16 | 5.2 | WH | starvation survival decreased, lifespan decreased | (Baugh and Sternberg, 2006; Kenyon *et al*, 1993; Munoz and Riddle, 2003) |
| MEF-2 | 6.6 | Other |  |  |
| ATF-7 | 6.6 | bZIP |  |  |
| LIN-14 | 6.0 | Other | lifespan increased | (Ewald *et al*, 2016) |
| JUN-1 | 5.3 | bZIP | lifespan decreased | (Khan *et al*, 2013) |
| HMG-1.1 | 7.1 | Other |  |  |
| NHR-25 | 9.1 | ZF - NHR |  |  |
| GRH-1 | 6.1 | Other | lifespan decreased | (Grigolon *et al*, 2022; Rozanov *et al*, 2020) |
| FKH-8 | 14.5 | WH |  |  |
| FOS-1 | -4.4 | bZIP |  |  |
| PHA-4 | -5.5 | WH | starvation survival decreased, lifespan decreased | (Panowski *et al*, 2007; Wu *et al*, 2018; Zhong *et al*, 2010) |
| Y75B8A.6 | -5.1 | Other |  |  |
| CEH-16 | -6.0 | HD |  |  |
| HMG-4 | 5.4 | Other | lifespan increased | (Tacutu *et al*, 2012) |
| ATF-2 | -5.2 | bZIP |  |  |
| CEH-48 | -7.2 | HD |  |  |
| LIN-32 | -5.2 | bHLH | lifespan increased | (Rozanov *et al*, 2020) |
| GSOX-1 | -4.3 |  |  |  |
| APTF-3 | -4.5 | Other |  |  |
| PAX-3 | 4.8 | HD |  |  |
| HLH-1 | 4.6 | bHLH |  |  |
| BCL-11 | 4.8 | ZF - C2H2 |  |  |
| DSC-1 | 4.9 |  |  |  |
| UNC-62 | 4.3 | HD | lifespan increased | (Catoire *et al*, 2011) |
| CTBP-1 | -4.8 | Other |  |  |
| HLH-11 | 5.9 | bHLH | lifespan increased | (Rozanov *et al*, 2020) |
| LIN-26 | -6.2 | ZF - C2H2 |  |  |
| DPL-1 | -6.9 | WH | starvation survival decreased | (Chen *et al*, 2025) |
| LIN-15B | -10.6 | AT Hook | starvation survival decreased | (Chen *et al*, 2025) |
| MADF-8 | 4.3 | Other |  |  |
| LSL-1 | 7.8 | ZF - C2H2 |  |  |
| ATTF-3 | 5.0 |  |  |  |
| NHR-20 | 4.6 | ZF - NHR |  |  |
| ZIP-2 | 6.2 | bZIP | starvation survival decreased | (Yan *et al*, 2025) |
| Y44A6D.3 | -6.0 | Other |  |  |
| ZNF-622 | -6.4 |  |  |  |
| PAT-9 | -5.4 | ZF - C2H2 |  |  |
| ATFS-1 | -5.0 |  | lifespan decreased | (Rauthan *et al*, 2013) |
| LIN-13 | -5.1 | ZF - C2H2 | lifespan decreased | (Knutson *et al*, 2016) |
| NHR-23 | -9.3 | ZF - NHR |  |  |
| HAM-2 | -7.1 | ZF - C2H2 |  |  |
| ZTF-27 | -4.4 | ZF - C2H2 |  |  |
| ATF-6 | -4.4 | bZIP |  |  |
| ZFH-2 | -7.5 | ZF - C2H2 |  |  |
| XBP-1 | -7.7 | bZIP | starvation recovery defective, lifespan decreased | (Chen *et al*, 2009; Henis-Korenblit *et al*, 2010; Roux *et al*, 2016) |
| SKN-1 | -5.6 | bZIP | starvation survival affected, lifespan decreased | (Bishop and Guarente, 2007; Lehrbach and Ruvkun, 2019; Paek *et al*, 2012) |
| C34B4.2 | -6.8 | WH |  |  |
| LET-607 | -10.0 | bZIP | lifespan increased | (He *et al*, 2021) |

Despite widespread tissue-specific effects of starvation, clustering revealed similarity among functionally related tissues. Many of the similarities seen at the cellular level (Fig. 2B) are evident at the tissue level as well (Fig. 4C). In addition, pharyngeal neurons (PharyngealNeuron), ciliated neurons (CiliatedNeuron), and all other neurons (Neuron) clustered together. Relatively small clusters of genes up or downregulated in many tissues are also evident (Fig. 4C). In fact, six (*M01H9.3*, *nmad-1*, *C16A3.6*, *cytb-5.2*, *F56H9.2*, and *T23G7.3*) and four (*Y22D7AL.10*, *inf-1*, *R06C1.4*, and *cyn-7*) genes were up and downregulated, respectively, in all eighteen tissues (Fig. 4B), suggesting a core starvation response across the animal. In summary, analysis of the starvation response at the cell and tissue levels reveals extensive variation among cells/tissues but also suggests shared effects among functionally related cells/tissues and some relatively rare effects shared across much of the animal.

### Functional consequences of starvation in different tissues

We used Gene Ontology (GO) term enrichments (Raudvere *et al*, 2019) to interrogate the function of genes affected by starvation in each tissue. We identified 53 GO terms enriched among DEGs in at least one tissue, and 33 of them were enriched in only one tissue (Fig. 5A-B, Sup. Data 5). The number of GO terms in each tissue ranges from two (GLR) to ten (Germline and SGP (somatic gonad precursor; Z1Z4)), with a median of five GO terms per tissue (Fig. 5C). These results suggest that starvation affects different ‘molecular functions’, ‘biological processes’, and ‘cellular components’ in different tissues, though sensitivity to detect GO term enrichments is related to the number of DEGs per tissue.

**Figure 5.**
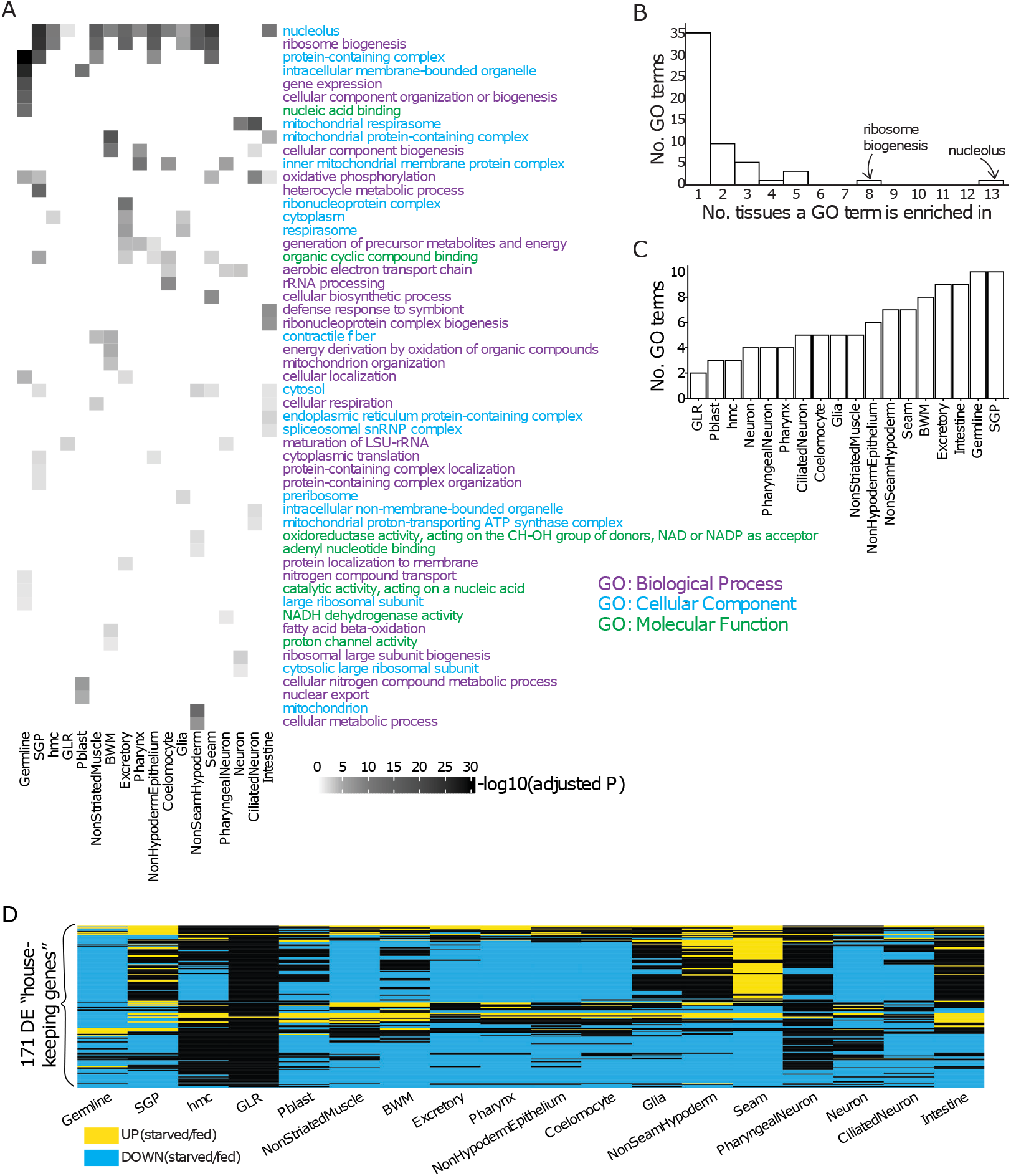
Tissue-specific enrichments of Gene Ontology (GO) terms based on differential expression. (A) A heatmap of GO terms enriched among DEGs in each tissue. The three GO term categories are colored differently. Shades of gray represent −log_10_(adjusted p-value) from GO term analysis. GO term analysis was performed with conditional probability being considered (Raudvere et al. 2019) and the background set being detected protein-coding genes in each tissue. Only ‘driver’ GO terms with an adjusted p-value of less than 0.05 were plotted. For detailed GO term analysis results, see Sup. Data 5. Column (tissue) order is the same as in Fig. 2E. (B) Number of GO terms enriched in a certain number of tissues. The two most widely enriched GO terms are labelled. (C) Number of GO terms enriched in each tissue plotted in rank order. (D) Heatmap showing up and downregulation of 171 ‘housekeeping genes’ (Ghaddar *et al*, 2023) differentially expressed in at least one tissue (all but one housekeeping gene were differentially expressed). Genes upregulated in starved compared to fed conditions are yellow; downregulation is blue. Heatmap rows were hierarchically clustered and reordered by optimal leaf ordering (OLO). Column (tissue) order is the same as in Figure 4C.

In addition to tissue-specific functional consequences of starvation, GO term enrichments suggest relatively universal effects of starvation on essential functions. The terms ‘ribosome biogenesis’ and ‘nucleolus’ are outliers for being enriched in eight and thirteen tissues, respectively (Fig. 5A-B), suggesting nutrient availability impacts translational capacity across the animal. These findings are consistent with bulk mRNA-seq, ribosome profiling, and proteomics experiments showing decreased expression of translation genes, translational activity, and ribosomal proteins during L1 arrest (Baugh *et al*, 2009; Maxwell *et al*, 2014; Stadler and Fire, 2013; Tuomaala *et al*, 2026).

The behavior of genes associated with translation prompted us to look at so-called ‘housekeeping’ genes, which were previously identified based on 1) being involved in essential functions (*e.g.*, translation and respiration), 2) having consistent expression across cell types in scRNA-seq analysis of adults, and 3) being conserved (Ghaddar *et al*, 2023). 171 of 172 housekeeping genes were differentially expressed in at least one tissue, with *tct-1/TPT1* being the exception (making it ideal for qPCR normalization), and in most cases they were downregulated (Fig. 5D). Notably, the housekeeping genes displayed largely consistent effects across tissues, consistent with their definition as housekeeping genes, though their expression is clearly affected by nutrient availability.

### Identification of candidate transcription factors mediating nutritional control of gene expression

We sought to elucidate transcriptional regulation of the starvation response. *Cel*EsT integrates *C. elegans* data from ChIP-seq, DNA-binding motifs, yeast one-hybrid screens, and conservation into a gene regulatory network designed to estimate the combined activity of transcription factors (TFs) from gene expression data (Perez, 2025). *Cel*EsT uses a multivariate model to identify the best combination of TF activities to account for the observed differences in gene expression between conditions. In principle, such an approach should be superior to univariate models that consider expression of a TF’s targets as a proxy for activity of that TF without considering potential contributions of other TFs. The gene regulatory network is unsigned (whether TFs function as activators or repressors is not considered), and we do not have enough information to interpret the significance of positive vs. negative TF activity values (Perez, 2025). We ran *Cel*EsT on each tissue after excluding TFs not detected in fed or starved larvae in that tissue (Sup. Data 6). The number of TFs with significant activity estimates ranged from one per tissue (GLR) to 37 (Intestine) with a median of 23 (Fig. 6A). 137 TFs had significant activity estimates in any tissue, with 67 (49%) in a single tissue (Fig. 6B; Sup. Data 6). Of the 70 TFs with significant activity estimates in at least two tissues, 53 (76%) had concordant effects across significant tissues. The number of TFs implicated in each tissue correlates with the proportion of detected genes that are differentially expressed in that tissue (p < 10^-4^; Sup. Fig. 7). Thus, like the results of differential expression analysis, TF activity analysis suggests a combination of common and tissue-specific gene regulatory mechanisms.

**Figure 6.**
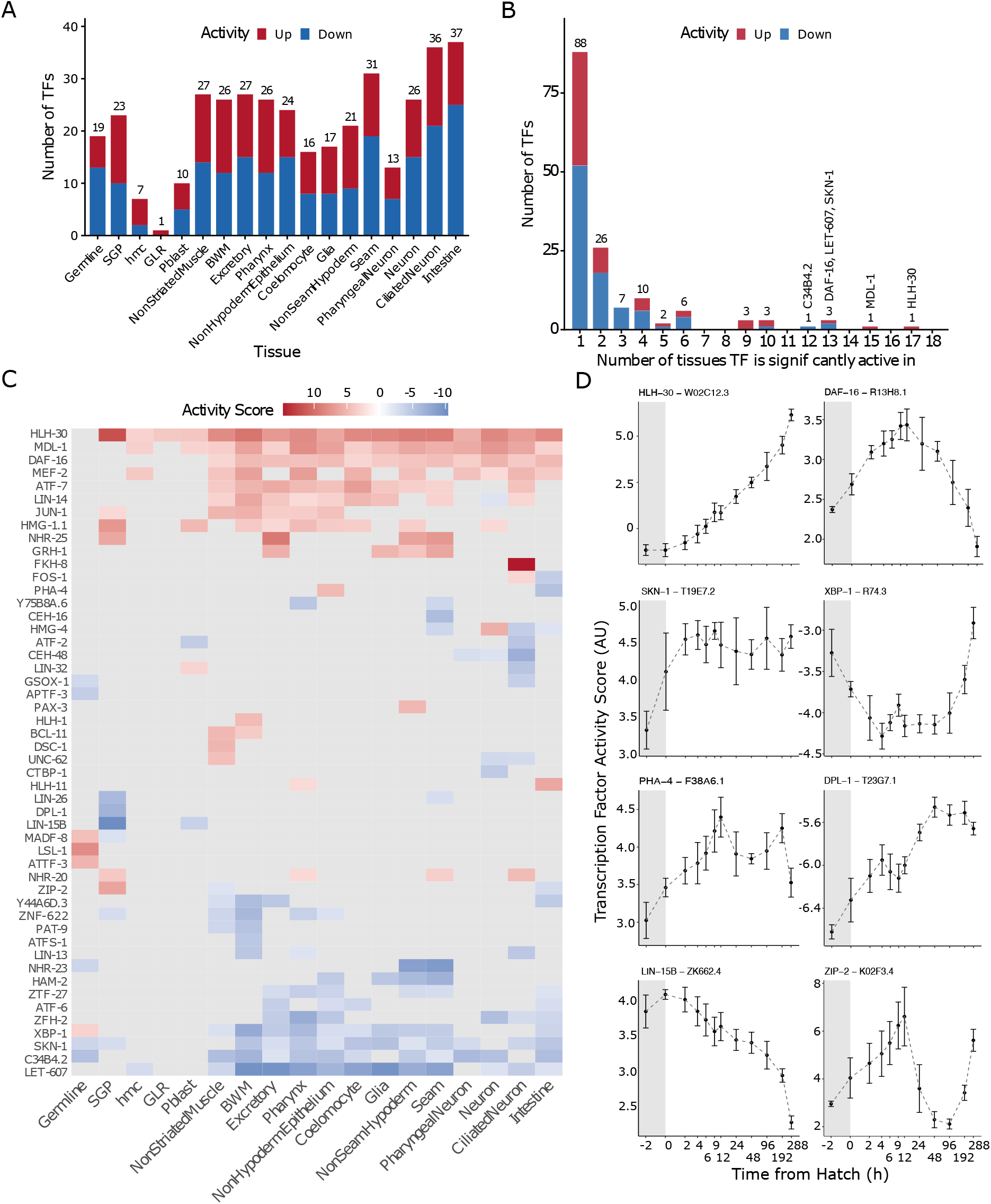
Inference of tissue-specific differential transcription factor (TF) activity in fed and starved L1 larvae. *Cel*EsT was used to infer differences in TF activity (Perez, 2025). *Cel*EsT generates positive and negative TF activity values, but interpretation of the significance of positive vs. negative values requires knowing whether each TF functions as an activator or repressor in every instance, and such interpretation is therefore not reliable. TFs not detected in each tissue were censored from analysis (see Sup. Data 6). (A) Bar chart plotting the number of TFs with significant activity scores (adjusted p < 0.05) in each tissue. (B) Bar chart plotting the number of tissues each TF is significant in. (C) Heatmap of TF activity scores after hierarchical clustering for the top 50 TFs based on their maximum absolute value tissue-specific activity scores. Column (tissue) order is the same as in Figure 4C. (D) Temporal dynamics of TF activity over 12 days of L1 starvation for eight TFs known to affect starvation survival (Table 1). *Cel*EsT was used to infer relative TF activity over time in a published bulk RNA-seq time series (Webster *et al*, 2022). The time series commenced approximately 2 hr before hatching (grey shading indicates pre-hatch) with relatively dense sampling early and sparser sampling later as the rate of change in expression dynamics decreases, and it was collected in four biological replicates. Time points sampled are indicated on the x-axis. Error bars reflect 95% confidence intervals based on activity estimates of four biological replicates per time point. Since expression levels were used rather than ratios (eg, starved/fed), activity scores are not comparable between genes. See also Sup. Fig. 7 and Table 1.

*Cel*EsT identified eight TFs that are known to promote the starvation response and support starvation survival. HLH-30/TFEB stands out as having a putative effect in 17 of 18 tissues (all but germline), and DAF-16/FoxO has a putative effect in 13 tissues (Fig. 6B, C, Table 1, and Sup. Data 6). *hlh-30* and *daf-16* both affect expression of thousands of genes during L1 arrest (Fisher *et al*, 2026; Munoz-Barrera *et al*, 2025), and both support starvation survival (Baugh and Sternberg, 2006; Munoz and Riddle, 2003; O’Rourke and Ruvkun, 2013), with *hlh-30* mutants dying faster than any other in the absence of food. Transgenic expression of *daf-16* in the intestine, epidermis, or nervous system is sufficient to partially rescue starvation survival of the null mutant (Kaplan *et al*, 2015), and DAF-16 activity is implicated in each of those tissues (Fig. 6C). SKN-1/Nrf has a putative effect in 13 tissues, and it is reported to contribute to the starvation response with complex effects on starvation survival (Paek *et al*, 2012). XBP-1/XBP1 has a putative effect in 11 tissues, and it is required for recovery from L1 arrest (Roux *et al*, 2016). PHA-4 has putative effects in two tissues (NonSeamHypoderm and Intestine), and it has thousands of binding sites in starved L1 larvae and promotes starvation survival (Zhong *et al*, 2010). DPL-1/Dp-1 is part of the DREAM complex which acts through the THAP domain-containing protein LIN-15B to repress germline gene expression promoting L1 starvation survival (Chen *et al*, 2025; Gal *et al*, 2021), and DPL-1 and LIN-15B are both implicated in the somatic gonad precursors (SGP). The bZIP TF ZIP-2 affects expression of over a thousand genes during L1 arrest to support starvation survival (Yan *et al*, 2025), and it is implicated in three tissues (Fig. 6C). These observations show that *Cel*Est identified TFs that are critical to the starvation response and starvation survival, suggesting that the other TFs identified are promising candidates.

*Cel*EsT also identified TFs that have not been implicated in starvation survival as candidate nutrient-dependent regulators of transcription. Sixteen of the top 50 TFs identified by *Cel*EsT (based on maximum absolute values of tissue-specific activity scores) affect adult lifespan (Table 1), supporting the hypothesis that they mediate nutritional control of gene expression. In addition, 10 of the top 50 TFs are from the bZIP TF family (p = 0.007; hypergeometric test). bZIP TFs have ancient eukaryotic functions in mediating transcriptional responses to various types of nutrient limitation in budding and fission yeast (Hinnebusch and Natarajan, 2002; Kanoh *et al*, 1996; Takeda *et al*, 1995) that are conserved and elaborated on in mammals (Neill and Masson, 2023). The bZIP and bHLH TF families were enriched across tissues, with each over-represented in 15 of 18 tissues (p = 0.002 and 0.04, respectively; Fisher’s combined probability test). The bHLH family includes HLH-30/TFEB, and like the bZIP family, bHLH family members mediate transcriptional responses to availability of various nutrients in budding and fission yeasts (Mercier *et al*, 2008; O’Neill *et al*, 1996) as well as mammals (Massari and Murre, 2000; Stine *et al*, 2015). These observations further suggest that *Cel*EsT identified promising TF candidates for nutrient-dependent transcriptional regulation in *C. elegans*.

We used *Cel*EsT to elucidate temporal dynamics of TF activity during L1 starvation to complement tissue-level analysis. A published bulk RNA-seq time series characterized gene expression from peri-hatch to 12 days of L1 starvation (Webster *et al*, 2022). We ran *Cel*EsT on variance-stabilized read counts for each of the 12 time points separately. Numerous TFs with inferred effects on their activities in the bulk time series were identified (Sup. Data 6), but we focus on the eight TFs identified from tissue-level analysis that are known to affect starvation survival (Table 1). These eight TFs displayed variable dynamics (Fig. 6D), suggesting that patterns of gene regulation change over time during starvation as the physiological consequences of starvation progress. For example, HLH-30 activity appears to climb steadily throughout the timeseries, while DAF-16 activity peaks at 12 h and falls thereafter. In contrast, SKN-1 activity appears to rise rapidly and then remain elevated, and LIN-15B activity progressively declines. XBP-1, PHA-4, and ZIP-2 display the most complex temporal dynamics, potentially reflecting different phases of the starvation response. Notably, *Cel*EsT analyzes steady-state transcript abundances without consideration of transcript decay rates, though transcript stability varies (Webster *et al*, 2022). Nonetheless, our starved scRNA-seq samples were collected 12 h after hatching, and temporal analysis of TF activities from the bulk RNA-seq time series suggests that the single-cell data captures a snapshot of a dynamic process unfolding.

### Widespread active transcription in starved, quiescent primordial germ cells

We were surprised to see 2,388 genes upregulated in starved PGCs (Germline; Fig. 4C; Sup. Data 3). PGC chromatin undergoes hyper-compaction (‘Stage II compaction’) during L1 arrest, resulting in global silencing of transcription (Belew *et al*, 2021; Webster *et al*, 2022). *daf-18/PTEN* represses PGC transcription during L1 arrest (Fry et al. 2021), and transcription has not been documented in starved PGCs. PGC chromatin undergoes decompaction upon feeding, which enables full-blown transcriptional activation (Wong et al. 2018). However, hyper-compaction does not happen instantaneously upon hatching in the absence of food, and starved PGCs in recently hatched L1 larvae are positive for phosphorylation of serine 2 of the C-terminal domain of RNAPII (pSer2) prior to hyper-compaction, suggesting transcriptional competence (Belew *et al*, 2021), though transcription has not been demonstrated.

We examined the data suggesting transcriptional activation in starved PGCs more carefully. We plotted differential expression for 161 ‘high-confidence’ germline genes (previously defined by meta-analysis of published results (Fry *et al*, 2021)). 113 of the high-confidence germline genes were differentially expressed, including 70 upregulated genes (Fig. 7A). These results support our annotation of this cell cluster as ‘germline’, and they suggest transcriptional activity in starved PGCs. We examined enrichment of KEGG pathways among germline DEGs (Kanehisa *et al*, 2025), and we found that six out of seven ‘DNA repair and replication’-related pathways were enriched (Sup. Data 7). Germ cells have greater capacity for DNA repair than somatic cells, and LIN-35/Rb and the DREAM complex repress germline gene expression in the soma thereby repressing DNA repair activity (Bujarrabal-Dueso *et al*, 2023). In addition, full-blown transcriptional activation in PGCs upon feeding causes DNA damage that is required for chromatin decompaction and must be repaired (Butuci *et al*, 2015; Wong *et al*, 2018). The enrichment of DNA repair-associated genes among germline DEGs suggests that the apparent germline differential expression is indicative of true regulation.

**Figure 7.**
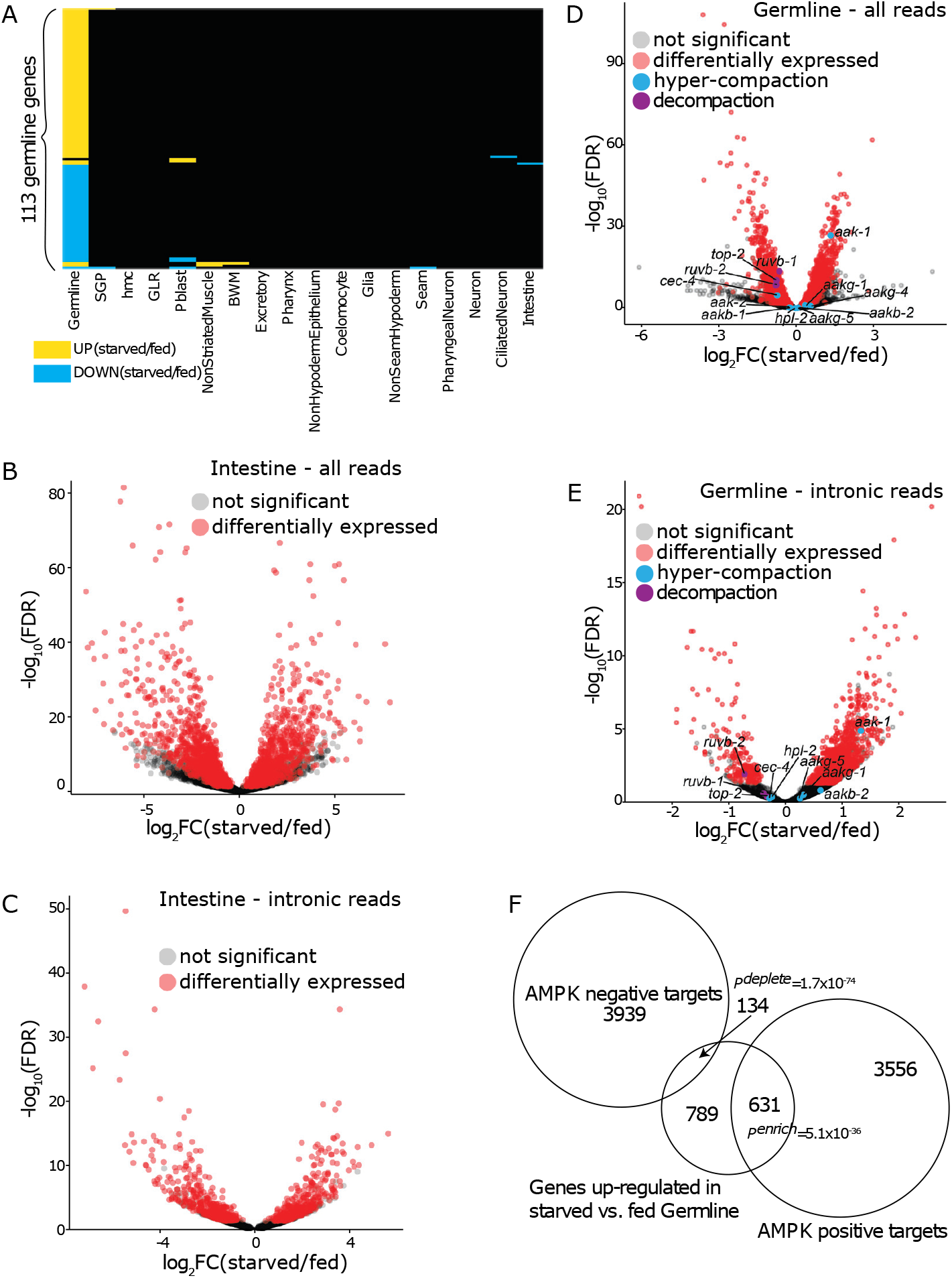
There is widespread active transcription in starved primordial germ cells. (A) Heatmap showing up and downregulation of ‘high-confidence’ germline genes (Fry *et al*, 2021) in different tissues. Genes upregulated in starved compared to fed conditions are yellow; downregulation is blue – degree of up or downregulation is not indicated. Column (tissue) order is the same as in Figure 4C. (B, C) Volcano plots showing −log_10_(FDR) vs. log_2_fold-change (FC) for all genes detected in the intestine. (D, E) Volcano plots showing - log_10_(FDR) vs. log_2_FC for all genes detected in the germline. Genes involved in primordial germ cell (PGC) chromatin hyper-compaction pathway (Belew *et al*, 2021) and genes involved in PGC chromatin decompaction pathway (Wong *et al*, 2018) are labelled. (B, D) DE analysis was done using all reads. (C, E) DE analysis was done using intronic reads only. (F) Overlap between genes upregulated in starved vs fed germline and AMPK positive and negative targets (El-Houjeiri *et al*, 2019). The background set for hypergeometric tests is all detected protein-coding genes in the germline. *P^enrich^* is enrichment p-value from the hypergeometric test. *P^delete^* is 1 minus *P^enrich^*. (B, D) See DataS4 for detailed DE analysis results using all reads in intestine and germline. (C, E) See DataS8 for detailed DE analysis results using intronic reads in intestine and germline. See also Sup. Fig. 8.

Apparent upregulation could result from transcription during starvation, or it could potentially result from transcript decay in fed larvae. We chose to analyze intronic reads as a proxy for transcriptional activity, since introns are only present in nascent transcripts. Introns can be detected since they are relatively A-T rich in *C. elegans* (facilitating cDNA priming with oligo-(dT)), but sensitivity of detection is limited. We first tested the approach in the intestine, a tissue for which we have no reason to doubt that there is transcription during starvation. 1,007 intestinal genes were upregulated in starvation based on the standard analysis of all reads (Fig. 7B), and 426 of these DEGs (42%) were upregulated based on analysis of intron reads alone (Fig. 7C; Sup. Data 8), with a significant correlation between fold-changes based on all reads and introns only (R = 0.96, p < 2.2 x 10^-16^, Sup. Fig. 8A). Given these reassuring results, we turned to the germline. Out of 1,554 upregulated germline genes based on all reads (Fig. 7D), 1,067 were upregulated (69%) based on intron reads alone (Fig. 7E; Sup. Data 8), and there was a significant correlation between fold-changes based on all reads and introns only (R = 0.93, p < 2.2 x 10^-16^, Sup. Fig. 8B). These results support the conclusion that many genes are transcribed in the PGCs during early L1 arrest.

### *aak-1/AMPK* is transcribed in primordial germ cells during early L1 arrest

Because PGC chromatin hyper-compaction occurs after larvae initially hatch in the absence of food (Belew *et al*, 2021), we wondered if any of the genes required for hyper-compaction or decompaction upon feeding were differentially expressed in the germline. The chromodomain protein-encoding gene *cec-4*, the heterochromatin protein 1 homolog-encoding gene *hpl-2/HP1*, and AMPK (which potentially includes components encoded by nine genes) are required for hyper-compaction (Belew *et al*, 2021), and *top-2/Topoisomerase II*, *ruvb-1/RUVB*, and *ruvb-2/RUVB* are required for decompaction (Wong *et al*, 2018). *top-2*, *ruvb-1*, and *ruvb-2* were each downregulated in starvation based on all reads (Fig. 7D), consistent with them being required for decompaction upon feeding. *ruvb-2* was also downregulated based on intronic reads, but *ruvb-1* and *top-2* were not (Fig. 7E; Sup. Data 8), presumably due to the lack of sensitivity in detecting intronic reads. Out of eleven genes potentially involved in hyper-compaction, only *aak-1* was upregulated during starvation (Fig. 7D), and it was also upregulated based on intronic reads alone (Fig. 7E; Sup. Data 8). These results suggest that *aak-1* is transcribed in the PGCs during early L1 arrest. Consistent with this, genes upregulated in starved vs. fed germline are significantly enriched for AMPK positive targets and significantly depleted for AMPK negative targets (Fig. 7F).

*aak-1* encodes one of two α subunits of AMPK, and *aak-2* encodes the other one. *aak-1* and *aak-2* share overlapping functions for most phenotypes (Fukuyama *et al*, 2012; Narbonne *et al*, 2017; Narbonne and Roy, 2006; Webster *et al*, 2017; Zheng *et al*, 2018), but loss of *aak-1* alone causes PGC divisions in starved L1s (Fukuyama *et al*, 2012). Simultaneous disruption of maternal and zygotic *aak-1* and *aak-2* with RNAi prevents hyper-compaction, but their individual effects were not investigated (Belew *et al*, 2021). These observations led us to hypothesize that AAK-1 is limiting for AMPK function in the PGCs of recently hatched L1s, such that transcription of *aak-1* in response to starvation is necessary for PGC chromatin hyper-compaction.

We used single molecule fluorescence *in situ* hybridization (smFISH) to confirm transcription of *aak-1* in starved PGCs. We designed 32 Quasar 670 (far red)-labelled oligonucleotide probes targeting multiple *aak-1* introns (Orjalo *et al*, 2011) in hopes of detecting nascent *aak-1* transcripts. We used an mCherry::H2B reporter expressed in the PGCs to label their chromatin, and we performed smFISH in wild-type and a new CRISPR *aak-1* deletion mutant (*aak-1(Δ)*). Given the bursty nature of transcription and the challenge of detecting nascent transcripts (Fukaya *et al*, 2016; Lee *et al*, 2017), we expected one or possibly two foci of Quasar signal co-localized with mCherry::H2B in a minority of PGC nuclei if *aak-1* is transcribed (Fig. 8A). Belew *et al*. reported that the PGCs of recently hatched (‘new L1’) larvae without food are positive for pSer2 and do not display Stage II hyper-compaction, and that pSer2 was lost and hyper-compaction was evident approximately 12 h later (‘starved L1s’). We performed smFISH in larvae that had recently hatched without food (12 h after bleach – ‘recently hatched L1’; Sup. Fig. 1) and in larvae that had experienced ∼12 h starvation (24 h after bleach – ‘12 h starved L1’). The later timepoint corresponds to our starved scRNA-seq samples that suggest accumulated *aak-1* mRNA as well as ongoing *aak-1* transcription (Fig. 7D-E), so we expected to detect *aak-1* introns after 12 h starvation and possibly in recently hatched L1s depending on the onset of transcription.

**Figure 8.**
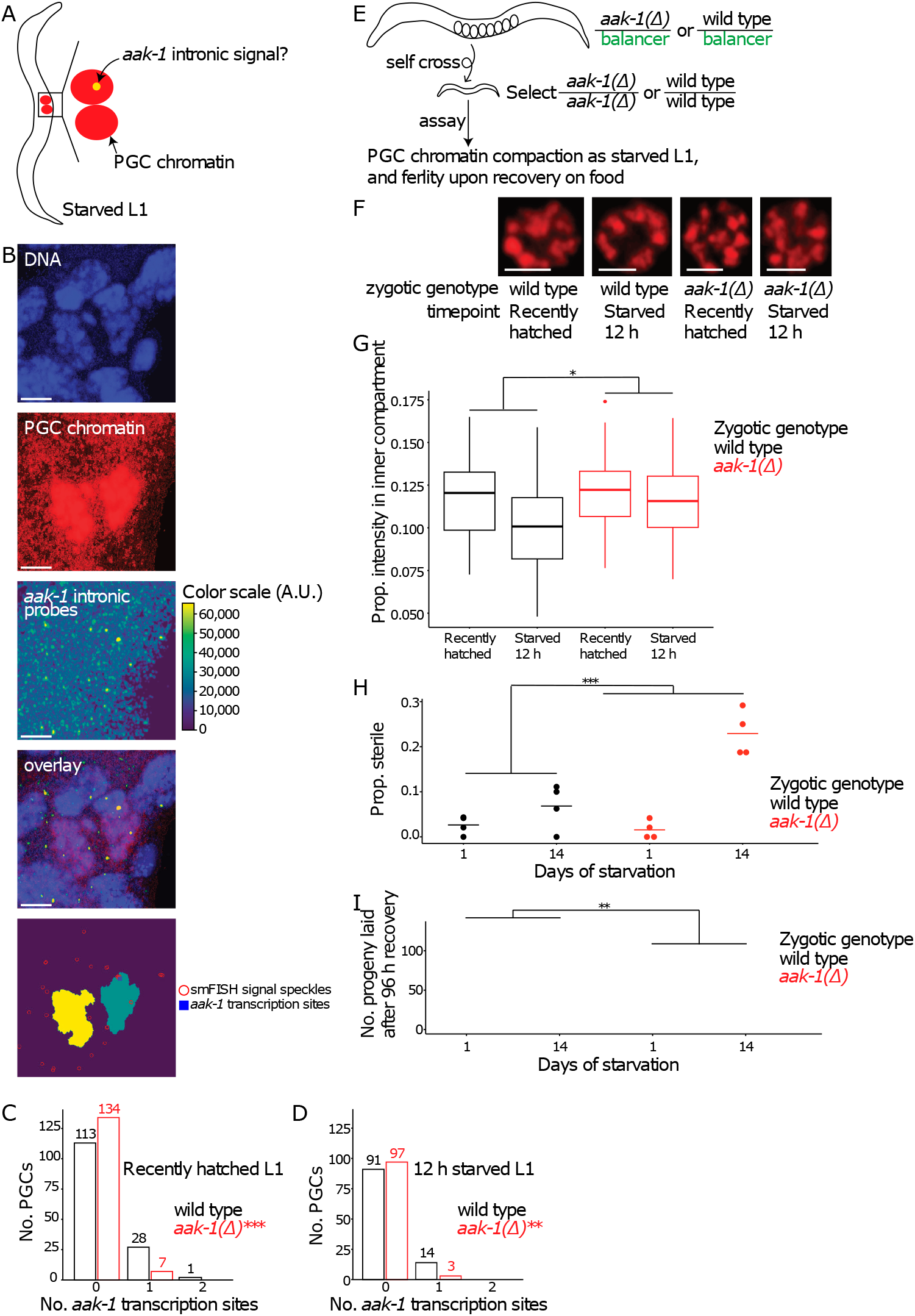
Zygotic transcription of *aak-1/AMPK* is required for chromatin hyper-compaction during starvation and fertility upon recovery. (A) Schematic of single-molecule fluorescent *in situ* hybridization (smFISH) experiment. (B) Representative smFISH images: (from top to bottom) DAPI-stained DNA, mCherry::H2B-labelled PGC chromatin, Quasar 670 (far red) labelled *aak-1/AMPK* intronic probe signal (signal intensity scale uses arbitrary units (A.U.)), overlay of these three channels, and computational reconstruction of images and identification of *aak-1/AMPK* transcription sites in PGC chromatin. (C, D) Number of PGCs with 0, 1, or 2 *aak-1/AMPK* transcription sites in PGC chromatin in wild-type vs *aak-1(Δ)* larvae. Fisher’s exact test was performed comparing *aak-1(Δ)* to wild type at each time point. (C) ‘Recently hatched L1’ was collected 12 h after bleach. Number of PGCs: 140 for wild type; 141 for *aak-1(Δ)* mutants. (D) ‘12 h starved L1’ was collected 24 h after bleach. Total number of PGCs: 105 for wild type; 100 for *aak-1(Δ)* mutants. (E) Schematic of experimental set-up to assay PGC chromatin hyper-compaction (F and G) and fertility upon feeding (H and I) in zygotic *aak-1(Δ)* mutants vs wild type. Balancer is pharyngeal GFP-marked *hT2*, and it is colored green in the schematic. (F) Representative PGC chromatin images showing different levels of compaction in recently hatched (12 h after bleach) and 12 h starved (24 h after bleach) wild-type and zygotic *aak-1(Δ)* larvae. (G) Proportion of PGC chromatin intensity in the ‘inner compartment’ (11.1% total area as defined in Belew *et al* in recently hatched and 12 h starved wild-type and zygotic *aak-1(Δ)* larvae. Number of PGCs: 70 for recently hatched wild type, 108 for 12 h starved wild type, 76 for recently hatched *aak-1(Δ)*, and 58 for 12 h starved *aak-1(Δ)*. Two-way ANOVA (formula: proportion intensity in the inner compartment ∼ genotype * duration of starvation) was performed, and the interaction-term p-value is presented. (H) Proportion of sterile wild-type and zygotic *aak-1(Δ)* worms that were recovered with food after 1 or 14 days of L1 starvation. Two-way ANOVA (formula: proportion sterile ∼ genotype * duration of starvation) was performed and the interaction-term p-value is presented. (I) Number of progeny laid in the first 96 hours of recovery by non-sterile wild-type and zygotic *aak-1(Δ)* worms that were recovered with food after 1 or 14 days of starvation. Linear mixed-effects (lme) model (dependent variable is number of progeny; fixed effect is genotype * duration of starvation; random effect is biological replicate) was fit to the data, and the fixed-effect interaction-term p-value is presented. (H, I) Four biological replicates were performed. Number of worms is 43 +/- 10 (mean +/- standard deviation). (B, F) Scale bar: 2 microns. (C, D,G-I) *P<0.05; **P<0.01; ***P<0.001. See also Sup. Fig. 9.

Our smFISH results suggest that *aak-1* is transcribed in starved PGCs. We collected Z-stack images, performed maximum intensity projection, and analyzed images using the Big-FISH pipeline (Imbert *et al*, 2022) (Fig. 8B). We chose analysis parameters such that the detection rates in *aak-1(Δ)* were ∼5% and ∼3% in recently hatched L1s and ∼12 h starved L1s, respectively (Fig. 8C-D). Our rationale was that these are acceptable false-positive rates for the negative control while maximizing detection sensitivity. In contrast, we detected *aak-1* transcription sites in 20% of PGCs in wild-type recently hatched L1s, and in 13% of PGCs in wild-type 12 h starved L1s (Fig. 8C-D), both of which are significantly different from *aak-1(Δ)*. These results suggest that starved PGCs transcribe *aak-1* at both timepoints. Notably, two sites of *aak-1* transcription were detected in only one wild-type PGC in a recently hatched L1, and the rates of detection were relatively low overall. Consistent with our results, relatively low rates of intron detection have been observed in other germline smFISH experiments (Lee *et al*, 2017). Low detection rate suggests transcription is sporadic such that not every *aak-1* locus is active at a moment in time, but it could also reflect lack of detection sensitivity.

### Transcription of *aak-1/AMPK* is required for chromatin hyper-compaction during starvation and fertility upon recovery

Having verified *aak-1* transcription in starved PGCs (Fig. 8A-D), we sought to test its functional relevance to test our hypothesis that it is required for chromatin hyper-compaction. AMPK is not required for Stage I PGC chromatin compaction during embryogenesis, but it is required for Stage II hyper-compaction in starved L1s (Belew *et al*, 2021). Belew *et al*. did not delineate maternal/zygotic requirements of AMPK, nor did they determine whether *aak-1* or *aak-2* alone was required for hyper-compaction. Our model predicts that zygotic *aak-1* is required. Homozygous *aak-1(Δ*) worms are viable, but we assayed the homozygous mutant self-progeny of *aak-1(Δ)* heterozygous mothers using a GFP-marked balancer chromosome (*hT2*) (Fig. 8E). Maternal *aak-1* was therefore functional, but zygotic *aak-1* was null. For a negative control, we analyzed homozygous wild-type self-progeny of wild type/*hT2* heterozygous mothers. We confirmed that wild-type PGC chromatin becomes hyper-compacted during L1 starvation (p = 1×10^-6^, Fig. 8F-G; Sup. Fig. 9A). *aak-1(Δ*) did not affect compaction in recently hatched L1s (p = 0.6), as expected given that AMPK is required for hyper-compaction, which should not have occurred yet (Belew *et al*, 2021). However, hyper-compaction was largely abolished in *aak-1(Δ)* 12 h starved L1s compared to recently hatched L1s (p = 0.08, Fig. 8F-G). Furthermore, there was a significant interaction between genotype and duration of starvation (Fig. 8G), supporting the conclusion that zygotic *aak-1* is required for hyper-compaction of PGC chromatin during early L1 arrest. Such functional requirement for zygotic *aak-1* reinforces the conclusion that *aak-1* is transcribed in starved PGCs.

We sought to extend our functional analysis of zygotic *aak-1* to reproductive success upon recovery from L1 arrest. The chromodomain-encoding gene *cec-4* is required for hyper-compaction and transcriptional silencing (Belew *et al*, 2021). Transcriptional silencing persists for days, likely indefinitely without feeding, and *cec-4* protects worms subjected to extended L1 arrest from sterility and reduced fecundity upon recovery (Webster *et al*, 2022). We hypothesized that since zygotic loss of *aak-1* disrupted PGC hyper-compaction in response to starvation, it would also increase sterility and decrease progeny production in adults that were subjected to extended L1 arrest. As for assaying hyper-compaction, we analyzed homozygous progeny of heterozygous *aak-1*mothers (Fig. 8E). Subjecting wild-type worms to 14 d L1 arrest increased sterility, as expected (Jobson *et al*, 2015; Webster *et al*, 2022), but sterility was approximately three-times more common with zygotic loss of *aak-1* (Fig. 8H). There was a significant interaction between genotype and duration of starvation on sterility, suggesting that zygotic *aak-1* protects worms from starvation-induced sterility. Likewise, the number of progeny produced in non-sterile wild-type worms within 96 h of recovery from L1 arrest decreased, as expected, but it decreased substantially more with zygotic loss of *aak-1* (Fig. 8I). There was a significant interaction between genotype and duration of starvation on early fecundity in non-sterile worms, suggesting that zygotic *aak-1* supports progeny production in fertile worms subjected to extended L1 arrest. This result suggests delayed onset of reproduction and/or reduced rate of progeny production with loss of zygotic *aak-1* and extended starvation, both of which, along with increased sterility (Fig. 8H), indicate decreased reproductive success. These results suggest that *aak-1* transcription in PGCs in response to starvation is required for chromatin hyper-compaction, germline transcriptional silencing, and reproductive success upon feeding.

## DISCUSSION

We report the first cellular gene expression atlas for a metazoan starvation response. We confirm that starvation has profound effects on gene expression, and we find that different cells and tissues respond to starvation in remarkably different ways. Genes related to translation are downregulated in many tissues, but differential expression of most genes is restricted to one or two tissues. We were surprised to find widespread transcription in primordial germ cells (PGCs) during starvation, as they were thought to be transcriptionally quiescent. We validated transcription of *aak-1/AMPK* in starved PGCs, and we showed that *aak-1/AMPK* transcription is necessary for PGC chromatin hyper-compaction and reproductive success after feeding. This work reveals remarkable tissue-specificity of the starvation response, and it shows that germline transcription is necessary to ultimately establish transcriptional silencing of the germline during starvation with implications for other quiescent cells.

### Anatomical complexity of the starvation response

Organisms have always confronted fluctuations in nutrient availability, and single-cell organisms have robust starvation responses, so one might expect the various cell types in an animal to autonomously mount a common response. However, anecdotally, reporter genes affected by nutrient availability are expressed in complex patterns (Baugh *et al*, 2009; Chen and Baugh, 2014; Hibshman *et al*, 2017), and with the evolution of multicellularity there was an elaborate specialization of cell function which may be expected to result in tissue-specific responses. Nonetheless, intuition suggests there must be common metabolic or gene regulatory adaptations to starvation that occur in most if not all tissues. Indeed, we found that genes related to translation are downregulated across the animal, along with other so-called ‘housekeeping genes’ (Fig. 5). However, multiple lines of evidence suggest that the starvation response is also tissue specific: 1) Most differential expression is specific to one or two cell types or a single tissue (Fig. 2, 4), 2) Most GO term enrichments are restricted to a single tissue (Fig. 5), and 3) Diverse combinations of TFs appear to promote the starvation response in different tissues (Fig. 6). The fact that we have more sensitivity to detect expression than differential expression likely contributes to each of these observations. Nonetheless, these results suggest that much of the starvation response is cell and tissue-specific.

Anatomical complexity of the starvation response implies an interface between developmental and environmental regulation of transcription. The response to other environmental stressors supports this hypothesis. *hlh-1/MyoD*, which promotes differentiation of muscle (Chen et al. 1994), regulates proteostasis in differentiated muscle cells by regulating tissue-specific chaperone expression (Nisaa and Ben-Zvi 2022). Likewise, the GATA factor gene *elt-2*, which promotes intestinal differentiation (Maduro and Rothman 2002), functions in the mature intestine to contribute to immunity, tissue-specific induction of zinc-responsive genes, and transient hypoxia-induced lifespan extension (Block *et al*, 2015; Roh *et al*, 2015; Schieber and Chandel, 2014). The *Drosophila* GATA factor gene *serpent* orchestrates tissue-specific outputs of the circadian oscillator (Meireles-Filho *et al*, 2014), supporting a general hypothesis about GATA factors and inducible gene regulation (Block and Shapira, 2015). We propose that developmental TFs with anatomically restricted expression interface with conditional, nutrient-responsive pathways to generate patterned responses to starvation. TFs identified by *Cel*EsT as potentially contributing to the starvation response in specific tissues (Fig. 6) are hypothetical candidates for such a function.

### Transcription and transcriptional silencing of the primordial germ cells during starvation-induced quiescence

We discovered widespread transcription in the two PGCs of L1 larvae that had just hatched in the absence of food (Fig. 2, 4, 7). We verified transcription of one of these genes, *aak-1/AMPK*, using smFISH (Fig. 8A-D). We were initially surprised to observe germline transcription, since we were under the impression that the PGCs do not initiate post-embryonic transcription until feeding. As discussed above, at hatching the PGCs have RNAPII pSer2, which suggests the capacity for transcription (Belew *et al*, 2021), but PGC transcription during L1 arrest has not been demonstrated. PGC chromatin is compacted at hatching, but it undergoes condensin-dependent hyper-compaction within the first ∼12 h of hatching without food (‘early L1 arrest’), and pSer2 is lost, indicative of transcriptional silencing (Belew *et al*, 2021). PGC chromatin undergoes decompaction upon feeding, and transcription commences (Wong *et al*, 2018). We previously reported that experimental degradation of the large subunit of RNAPII, AMA-1, specifically in the PGCs during L1 arrest does not affect transcript absolute abundance, confirming global transcriptional silencing in the PGCs (Webster *et al*, 2022). However, that experiment relied on the auxin-inducible degron (AID) system to degrade AMA-1, and we initiated exposure of worms to auxin ∼36 h after hatching without food; *i.e.*, after hyper-compaction occurs. We conclude that PGCs are transcriptionally active peri-hatching, but that transcription is silenced once the chromatin becomes hyper-compacted.

Our functional analysis of *aak-1/AMPK* suggests that PGC transcription in response to starvation is required for chromatin hyper-compaction and transcriptional silencing (Fig. 8E-I). Belew *et al*. showed that AMPK is required for hyper-compaction, but they did not determine if *aak-1* or *aak-2* alone was required, nor did they distinguish between maternal and zygotic requirements. There is no growth during *C. elegans* embryogenesis, and maternal products of many genes are sufficient for function in L1 larvae. However, we showed that zygotic *aak-1* is required for PGC chromatin hyper-compaction and reproductive success upon recovery (Fig. 8E-I). These results better define the genetic requirements for PGC hyper-compaction while demonstrating physiological significance. Moreover, these results are significant for demonstrating that transcription is necessary to establish transcriptional silencing. Critically, hyper-compaction is not a consequence of transcriptional silencing in this case. Inhibition of transcription in mammalian cells causes chromatin compaction (Neguembor *et al*, 2021), and we found that post-embryonic degradation of AMA-1 in the PGCs with AID caused severe chromatin compaction that appears distinct from hyper-compaction in wild-type worms (Sup. Fig. 9B, C). To the contrary, zygotic requirement of *aak-1/AMPK* for hyper-compaction shows that transcription is required for transcriptional silencing in starved PGCs.

Transcriptional silencing is a hallmark of PGCs and quiescent stem cells (Cheung and Rando, 2013; Seki *et al*, 2007; Seydoux *et al*, 1996; Shirae-Kurabayashi *et al*, 2011; Wessel *et al*, 2014), and chromatin compaction appears to be an ancient, conserved starvation response. Quiescent yeast cells undergo condensin-dependent chromatin compaction, silencing transcription, and this behavior is conserved in human fibroblasts (Swygert *et al*, 2019). Ribosomal DNA undergoes selective condensin-dependent chromatin condensation in response to glucose starvation in yeast (Xue and Acar, 2018). Likewise, chromatin and the nucleolus are reorganized in response to starvation in *C. elegans* intestinal cells (Al-Refaie *et al*, 2024). These findings resonate with PGC chromatin hyper-compaction during starvation-induced developmental arrest in *C. elegans*, suggesting a conserved role for AMPK in governing chromatin compaction during regulation of growth and quiescence.

## MATERIALS AND METHODS

### Strains used in this study

Wild type N2 is from the Sternberg Lab collection.

GS8729 *arSi12 [mex-5p::ERK::KTR::GFP(smu-1 introns)::T2A::mCherry::his-11::tbb-2 3’UTR] I* is from the Caenorhabditis Genetics Center (CGC).

GC1171 *naSi2[pGC550(Pmex-5 < mCherry::H2B::nos-2 3’ UTR < GFP::H2B::nos-2 3’ UTR - unc-119(+))] II; unc-119(ed3) III* is from the Hubbard Lab at New York University (Roy, Hubbard 2018).

### Strains generated in this study

PHX9618 *aak-1(syb9618) III* (‘*aak-1(Δ)*’) was generated by SunyBiotech. 5’ to 3’ sequence of *aak-1(syb9618)* is below, following this order:

upstream sequence/FIRST EXON OF *aak-1*[<u>3,778 bp deletion]</u>LAST EXON OF *aak-1*/downstream sequence

tgacctcctcagaccctatatattatttttgtttcgcggcatagttttcggaacttactgaaaacatttaattcttctaggaaaaca agccggatttcttggttaaacttttttaaaaagcttagttttgaaaatccctttgaaattttacccgccgccgagcctaagccaa agttttctccaaattttcaagtaaattttcaagccgatttttttttaatttttcgatttcacactacctgtcattaactcccaccgttt aacttttttaaaatccagaataataatgttttatttagggtatggtagtaccaataggttcacaacatcgtcaaaaaaatATGC CTC[<u>3,778 bp deletion</u>]CATCATGCAGGCTTTGTTAGCTGAATAAgaacataattttatgtttctatct

gtataatgatatgtatttttcgtttctttttcaaaattcatcttctcccggtttcacttttgccccccttcattcccctcagatgccttt tttctcttcaaagcatgtacttaaattggacgtctttgtttgttttttttttgttttattattctatttctgattccaattcaacaaggtagt gttatttgaaaaataataaaaatgattcaagaaattgaagaaattggccgtggagttggacccattgtagacaaaactcggtc gagccactgaaaattgtattatttctaaagaaaacaagaatttcagaacttacagccaatggagcat

LRB596 *+/hT2 [bli-4(e937) let-?(q782) qIs48] (I;III)*

LRB663 *arSi12[mex-5p::ERK::KTR::GFP(smu-1 introns)::T2A::mCherry::his-11::tbb-2 3’UTR] I; ieSi64 [gld-1p::TIR1::mRuby::gld-1 3’UTR + Cbr-unc-119(+)] II; ama-1(syb1513 [ama-1::GSGGGG::degron::TEV(ENLYFQSGK)::3xFLAG]) IV*

LRB686 *aak-1(syb9618) III/hT2 [bli-4(e937) let-?(q782) qIs48] (I;III)*

LRB687 *naSi2[pGC550(Pmex-5 < mCherry::H2B::nos-2 3’ UTR < GFP::H2B::nos-2 3’ UTR - unc-119(+))] II; aak-1(syb9618 deletion) III/hT2 [bli-4(e937) let-?(q782) qIs48] (I;III)*

LRB688 *+/hT2 [bli-4(e937) let-?(q782) qIs48] (I;III); naSi2[pGC550(Pmex-5 < mCherry::H2B::nos-2 3’ UTR < GFP::H2B::nos-2 3’ UTR - unc-119(+))] II*

### *C. elegans* maintenance

All strains assayed in this study were maintained with *E. coli* OP50 on nematode growth medium (NGM) plates and were well-fed for at least three generations before being used in experiments. Worms were cultured and starved at 20°C. Unless otherwise noted, all procedures were done at 20°C.

### Harvesting fed and starved samples

A synchronous population of gravid wild-type *C. elegans* (N2), grown on *E. coli* OP50-seeded plates, was treated with sodium hypochlorite solution to isolate embryos (‘bleach’; (Hibshman *et al*, 2021)). The collected embryos were resuspended in 10 mL of virgin S-basal (no ethanol or cholesterol) at a density of 5 eggs/µL to establish the starved culture, which was incubated at 20°C in a shaking incubator at 180 rpm.

A separate set of embryos was collected six hours after the starved culture was set up (∼6 h after hatching), using the same sodium hypochlorite treatment. These embryos were resuspended in 10 mL of S-complete buffer supplemented with 25 mg/mL (1x) *E. coli* HB101 at a density of 5 eggs/µL to establish the fed culture. Like the starved culture, the fed culture was incubated at 20°C in the same shaking incubator at 180 rpm. The starved culture was incubated for 24 hours, while the fed culture was incubated for 18 hours to allow approximately 12 hours of L1 starvation and six hours of L1 larval feeding, respectively (Sup. Fig. 1).

### Hatching curve for single-cell RNA-seq

Embryos were collected and cultured for fed and starved samples in the same way as in Harvesting fed and starved samples. Starting 10 h after hypochlorite treatment (‘bleach’), a 100 μL aliquot was sampled from the culture every hour, and the numbers of hatched and unhatched embryos were recorded. Proportion hatched was calculated as the number of hatched embryos divided by the total number of sampled embryos. The last time point of scoring was 18 h after bleach.

### Single-cell dissociation

Following incubation, fed worms were transferred to a 15 mL conical tube, washed three times with virgin S-basal to remove residual HB101 bacteria, and resuspended in 10 mL of virgin S-basal. The starved culture was collected at the same time and was similarly transferred to a separate 15 mL conical tube. Both tubes were centrifuged at 3,000 rpm for 1 minute to pellet the worms. The pellets were then transferred to individual 1.5 mL Eppendorf LoBind tubes (Cat No. 0030108523) using glass Pasteur pipettes and washed once with 1 mL of egg buffer (118 mM NaCl, 48 mM KCl, 2 mM CaCl_2_, 2 mM MgCl_2_, and 25 mM HEPES).

To disrupt the cuticle, 500 µL of freshly prepared SDS–dithiothreitol (DTT) solution (20 mM HEPES (pH 8.0), 0.25% SDS, 200 mM DTT, and 3% sucrose) was added directly to the worm pellet and incubated at room temperature (RT) in a ThermoMixer at 800 rpm for 2.5 minutes. Immediately after SDS-DTT treatment, 800 µL of egg buffer was added, and the worms were centrifuged at 21,100 x g for 30 s. The supernatant was aspirated, and the pellet was washed five times with 1 mL of egg buffer before transferring the worms to clean 1.5 mL tubes.

For enzymatic dissociation, 125 µL of freshly prepared pronase solution (Sigma P8811, 20 mg/mL in sterile egg buffer) was added to the SDS-DTT-treated pellet and incubated at RT in a ThermoMixer at 1,200 rpm. Additionally, the reaction was manually mixed by pipetting the entire volume up and down ∼60 times every three minutes to aid tissue dissociation. The dissociation process was monitored periodically by placing a 1–2 µL drop of the reaction mixture onto a slide and observing under a benchtop microscope.

After 18–20 minutes of pronase treatment, when single-cell dissociation was visually confirmed, the reaction was quenched by adding 1 mL of ice-cold L15-10% FBS (Invitrogen 21083027, Gibco A5256701) cell culture media. The cell suspension was left on ice for ∼10–15 minutes to allow unlysed worms and larger undigested tissue fragments to settle by gravity. The top 80% of the supernatant was transferred to a clean tube and centrifuged at 4°C for 10 minutes at 800 rpm. The cell pellet was washed twice with 1 mL of L15-10% FBS media and resuspended in 1 mL of culture media. Cell concentration was determined by pipetting 10 µL of the resuspended solution onto a hemocytometer and counting the cells on a compound microscope.

### scRNA-seq library preparation

Single-cell capture and library preparation were performed following the Chromium Next GEM 3′ Single Cell v3.1 protocol (10x Genomics 1000120, 1000268). For each channel, approximately 17,000 cells were mixed with the reverse transcriptase reaction solution and immediately loaded onto the capture chip to minimize exposure time to the reverse transcription cocktail. Final libraries were constructed using 25% of the cDNA volume from the reverse transcription reaction and 12 PCR cycles. Seven biological replicates were prepared for each culture condition (fed and starved), generating a total of 16 barcoded libraries (one biological replicate has two technical replicates). The pooled libraries were sequenced on a NovaSeq X Plus S-Prime flow cell using 10X Genomics default run parameters.

### scRNA-seq read mapping

Reads were mapped to the *C. elegans* reference transcriptome WS286. Due to the possibility that 3’ untranslated region (UTR) annotation in the reference transcriptome may be too short (Packer et al. 2019), we dynamically extended the 3’ UTR of each gene to its optimal length, enabling additional mapping of reads to the 3’ UTR of genes. We generated eight versions of gene annotations based on WS286 annotation, with 3’ UTR in each version elongated by 50, 100, 150, 200, 250, 300, 400 and 500 base pairs (bps), respectively. Elongation was only performed for genes whose 3’ end does not spill into another gene and was terminated before encountering the adjacent. Eight 3’ UTR extended genome annotation versions were used to map reads using scKB (Hsu *et al*, 2022). The number of reads mapped for each library are in Sup. Data 1.

For each of our 17 sequencing libraries, we generated read counts per gene per extended genome annotation. Next, we calculated the total count per million (CPM) for each gene, combining its CPMs in fed and starved samples, for each replicate for each extended genome annotation. We didn’t want to extend 3’ UTR for marginally detected genes, and therefore only set out to determine the optimal 3’ UTR extension for genes whose fed plus starved CPM in a replicate is at least 1. To determine the optimal 3’ UTR extension for each gene, we applied a dynamic ‘voting’ strategy. More specifically, for each replicate, we recorded the 3’ UTR extension length for each gene that resulted in the highest CPM, and we determined the final ‘optimal’ 3’ UTR extension length as the one identified by the most replicates, requiring a consensus of at least two replicates. If the optimal 3’ UTR extension increases the gene’s CPM by at least 5%, compared to using the original genome annotation, we kept that optimal 3’ UTR for that gene. Lastly, with the 3’ UTR optimally extended genome annotation, we re-mapped genes in all samples using scKB (Hsu *et al*, 2022), and we generated a gene-by-cell raw count matrix for spliced, unspliced, and all reads for each sample. These matrices were used in *scRNA-seq data processing*.

### scRNA-seq data processing

In the gene-by-cell matrices generated in *scRNA-seq read mapping*, genes with no counts in any cell were removed. Cells were filtered based on the following criteria: 1) We flagged ambient RNA using the COPILOT pipeline (Hsu *et al*, 2022); 2) Low-quality cells (presumably dying) were identified based on the enrichment of mitochondrial gene expression (>=20% of the total UMI counts; consistent with (Taylor *et al*, 2021)) (Sup. Fig. 2); 3) The expression profile of ‘dying cells’ was entered into COPILOT for it to iteratively and adaptively classify cells into high- and low-quality groups, and low-quality cells were censored; 4) To further remove outliers, cells with more than 38,041 unique molecular indices (UMI; top 1% UMI counts in high-quality cells) were censored; and 5) We used the COPILOT pipeline to remove multiplets using the DoubletFinder algorithm (Hsu *et al*, 2022; McGinnis *et al*, 2019). After all this censoring, we have a filtered version of gene-by-cell count matrix for unspliced, spliced, and all reads for each sample. Those filtered matrices were used in *Differential expression (DE) analysis*.

### Differential expression (DE) analysis

Filtered gene-by-cell matrices generated in *scRNA-seq data processing* were aggregated to create gene-by-rep count matrices for unspliced and all reads. These matrices were entered into DESeq2 (Love *et al*, 2014). DE analysis was performed following the default DESeq2 pipeline with *ashr* being the log_2_FoldChange shrinkage method (see DESeq2 vignette for scRNA-seq analysis at https://www.bioconductor.org/packages/release/bioc/vignettes/DESeq2/inst/doc/DESeq2.html). Percentage of cells expressing each gene in starved and fed conditions, respectively, was calculated using Seurat’s *FoldChange* function (Butler et al. 2018). Criteria for differentially expressed genes (DEGs) between starved and fed conditions: 1) percentage of cell expressing this gene is at least 0.1 in either starved or fed conditions; 2) adjusted p-value of DE analysis is 0.05. For pseudobulk DE analysis, we aggregated all cells that pass quality-control filters (see scRNA-seq data processing for details) from each replicate in fed and starved conditions, separately. We then performed DE analysis using DESeq2 as described above to get the ‘pseudobulk’ DE expression profile.

### Hierarchical clustering

In all heatmaps where rows are genes and columns are cell types or tissues, each gene’s log_10_ fold-change (starved/fed) across different cell types or tissues was used for hierarchical clustering using the *hclust* function in R. Pearson distance and the complete linkage method were used. When Optimal Leaf Ordering (OLO) was performed, it was done using the *reorder* function in the seriation R package, with the method parameter set to ‘OLO.’ In Figure 5A, rows were hierarchically clustered using −log_10_(adjusted P) across tissues, and Euclidean distance and the complete linkage method were used.

### *Cel*EsT inference of transcription factor activity

#### Tissue level analysis

Wald statistics were calculated for each gene as the ratio of its log_2_ fold-change to the corresponding standard error, which were obtained from the DE analysis. These statistics were used as input for TF activity estimation using the decoupleR package in R and the orth*Cel*EsT gene regulatory network (GRN) downloaded from github.com/IBMB-MFP/CelEsT-app/. Prior to analysis, the GRN was filtered to retain only TFs detected in at least 10% of cells in either the fed or starved condition for each tissue. TF-family enrichment among significant TFs was assessed separately for each tissue using Fisher’s exact test. For each tissue, the background set consisted of all TFs detected in that tissue according to the same 10% expression threshold. P-values were adjusted for multiple testing across TF families using the Benjamini–Hochberg (BH) procedure. To evaluate whether enrichment was consistent across tissues, tissue-specific P-values for each TF family were combined using Fisher’s combined probability test, followed by BH correction across TF families. The direction of enrichment was assessed using the odds ratios from tissue-specific tests. TF-family enrichment was also assessed among the 50 TFs with the maximum absolute activity scores across tissues. Fisher’s exact tests were performed using a combined background comprising the union of the tissue-specific, detection-filtered TFs, with duplicate TFs removed. P-values from these tests were adjusted using the BH procedure.

#### Temporal analysis of bulk RNA-seq time series

Raw gene-level read counts from a previously published whole animal (bulk) RNA-seq L1 timecourse of L1 starvation from 2 h prior to hatch up to 12 d post-hatch (Webster *et al*, 2022) were downloaded from the supplementary file of accession GSE173656 in the Gene Expression Omnibus (GEO). Raw counts were normalized with a variance-stabilizing transformation using the *vst* function of DESeq2 in R. VST-normalized expression levels for each sample were used as input for TF activity estimation using a multivariate linear model (mlm) using the decoupleR package in R with the *Cel*EsT v1.1 gene regulatory network (GRN; (Perez, 2025)) downloaded from github.com/IBMB-MFP/CelEsT-app/. Before applying the mlm, the *Cel*EsT v1.1 GRN was filtered to remove TFs which were not detected with transcripts per million (TPM) > 1 in at least 4 samples – this led to the exclusion of 28 TFs (Sup. Data 6). As TF activity estimates were based on sample gene expression rather than differential expression analysis, the overall level of TF activity – the arbitrary units corresponding to the TF mlm score – cannot be compared across TFs. To examine TF activity dynamics in Fig. 6D, we plotted the mean mlm score for the four replicates of each time point with error bars corresponding to a 95% confidence interval calculated by the mean ± 1.96 × the standard error of the replicates. The x axis (length of L1 starvation) was transformed by a pseudo-log_2_ transformation to better visualize across the whole timecourse, which was more densely sampled in early starvation.

### smFISH experiment imaging and analysis

Worm samples were prepared for smFISH identical to the way they were prepared for scRNA-seq, except they were collected 12 h (‘recently hatched L1’) and 24 h (‘12 h starved L1’) after hypochlorite treatment. smFISH was performed following a previously published protocol (Lee *et al*, 2017) with the working stock concentration of smFISH probes being 5 μM and the final probe concentration in hybridization buffer being 50 nM (1:100 dilution of probes). 32 custom Stellaris smFISH Probes (each 22 nt long) were designed against *aak-1* introns (Biosearch Technologies, Inc., Petaluma, CA) and their sequences are in Sup. Table 1. After 24 h curing, fixed samples were imaged on a Zeiss LSM 880 Airyscan with a 63x oil immersion objective. Z-stacks were taken for each of mCherry, DAPI, and far-red channels at 0.21-micron intervals. Images were processed using ImageJ (Schneider *et al*, 2012) and Python following the Big-FISH pipeline (Imbert *et al*, 2022) (https://github.com/jc271828/scRNA-seq).

### Chromatin compaction imaging and analysis

Worm samples were prepared for imaging identical to the way they were prepared for scRNA-seq, except they were collected 12 h (‘recently hatched L1’) and 24 h (‘12 h starved L1’) after hypochlorite treatment (for Fig. 8E-F) or 12 h and 36 h after hypochlorite treatment (Sup. Fig. 9A). For the experiment that used auxin-induced degradation (AID) to degrade AMA-1 (Sup. Fig 9C), sample condition and auxin addition time point are in Sup. Fig. 8B. Because indole-3-acetic acid (IAA) auxin is dissolved in ethanol, we added 0.25% ethanol (the solvent) as a control to adding 1 mM auxin. IAA auxin addition concentration and the control ethanol addition concentration follow a previous paper using the same alleles (Webster *et al*, 2022). Live imaging was done on Zeiss LSM 880 Airyscan with a 63x oil-immersion objective. Z-stacks were taken at 0.21-micron intervals. A custom automated pipeline was designed to unbiasedly quantify chromatin compaction (https://github.com/jc271828/scRNA-seq). For each z-stack, an ImageJ script extracts PGC chromatin based on germline-expressed mCherry::H2B from each slice. Output of this step is a region of interest (ROI) mask TIFF and a segmented PGC chromatin TIFF for each slice of the stack. These pairs of TIFFs went into a custom Python program, and PGC chromatin was fit with a bounding box for each slice. The centroid of the PGC chromatin was defined as the center of the bounding box. A circle with area 11.1% of the ROI area (11.1% was first defined in (Belew *et al*, 2021)) was drawn and centered in the bounding box, and integrated signal intensity in this circle was defined as ‘intensity in inner compartment.’ This value was divided by total integrated intensity within the ROI to get ‘Prop. intensity in inner compartment.’ Because every imaged PGC nucleus was captured by a z-stack, the final step of the pipeline was to pick the slice from each stack with the largest ROI for quantification. Image settings were the same for all samples.

## Data availability

Raw scRNA-seq data files generated for this study have been deposited in the NCBI GEO SRA database (http://www.ncbi.nlm.nih.gov/sra) under accession number GSE296325 (access token ihebouiwlrotlyb). Code to reproduce all results presented in this study can be found at https://github.com/lrbaugh/scRNA-seq.

## ACKNOWLEDGEMENTS

We would like to thank the late Philip Benfey for providing access to the 10x Genomic Chromium instrument for scRNA-seq, Che-Wei Hsu and Trevor Nolan for their assistance with scRNA-seq techniques and analysis, and Seth Taylor and David Miller for helpful discussions. We would like to thank ChangHwan Lee for help with smFISH protocols. We would also like to thank WormBase and the Alliance of Genome Resources (Alliance of Genome Resources, 2024; Sternberg *et al*, 2024). This work was funded by the National Institutes of Health (R01GM117408, R01GM143159, and R35GM156356 awarded to L.R.B.).

## Author contributions

Conceptualization: L.R.B.; Data curation: J.C. and S.S.; Formal analysis: J.C., S.S. and M.F.P.; Funding acquisition: L.R.B.; Investigation: J.C., R.C. and J.R.V.; Methodology: L.R.B., M.F.P. and J.C.; Project administration: L.R.B.; Software: M.F.P; Resources: L.R.B.; Supervision: L.R.B.; Validation: J.C. and J.R.V.; Visualization: J.C., S.S. and M.F.P.; Writing – original draft: L.R.B.; Writing – review & editing: L.R.B, S.S. and M.F.P.

## SUPPLEMENTARY MATERIALS

**Sup. Figure 1.**
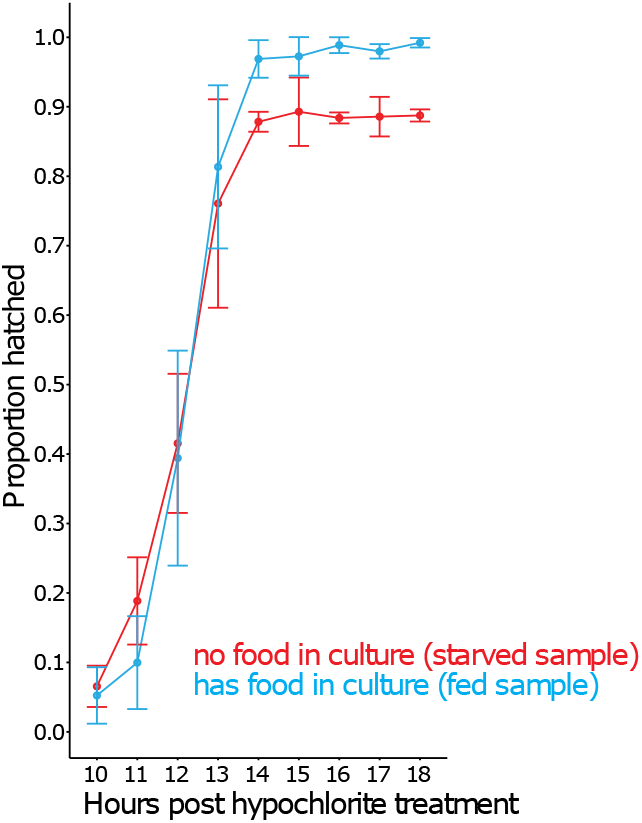
Proportion of hatched embryos (hatching efficiency) was measured between 10 and 18 h after hypochlorite treatment (‘bleach’). Related to Fig. 1A. Three biological replicates were performed. Dots are average hatching efficiency across replicates. Error bars represent standard deviation (SD). At ∼12 h after bleach, both starved and fed samples reached approximately 50% hatching. Therefore, 12 h after bleach was designated as the time point for hatching. Number of examined embryos (hatched and unhatched) is 96 +/- 15 (mean +/- SD).

**Sup. Figure 2.**
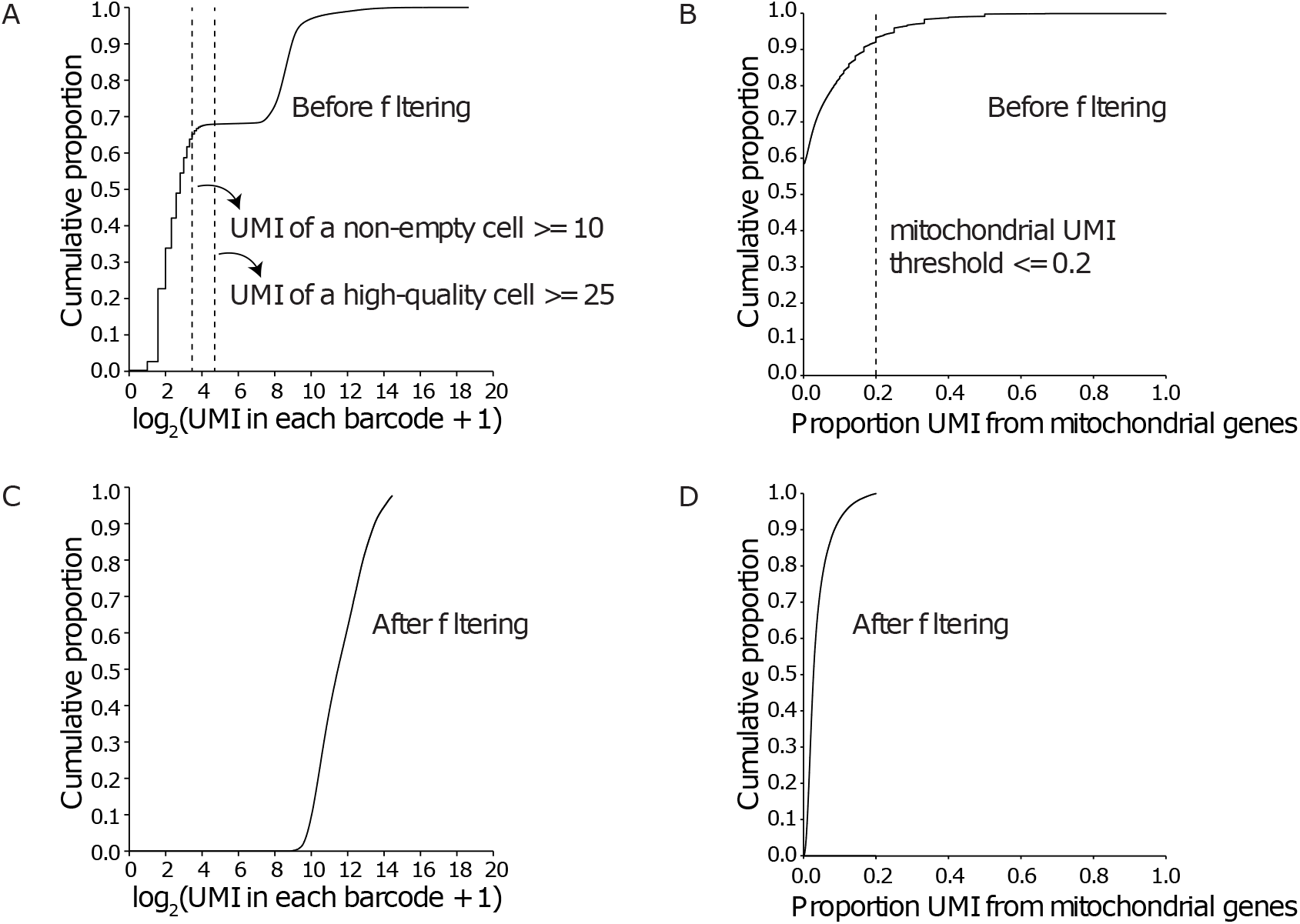
Quality control for single-cell RNA-seq samples. Related to Fig. 1. (A) Plot showing the cumulative distribution of log_2_-transformed UMI counts in each barcode (cell) before filtering. Filtering used at least 10 UMI and at least 25 UMI as the cutoff for non-empty cells and high-quality cells, respectively. (B) Plot showing the cumulative distribution of the proportion of UMI coming from mitochondrial genes in each barcode (cell) before filtering. Filtering used 0.2 as the cutoff for healthy cells (Taylor *et al*, 2021) (C) Plot showing the cumulative distribution of log_2_-transformed UMI counts in each barcode (cell) after filtering. (D) Plot showing the cumulative distribution of the proportion of UMI coming from mitochondrial genes in each barcode (cell) after filtering.

**Sup. Figure 3.**
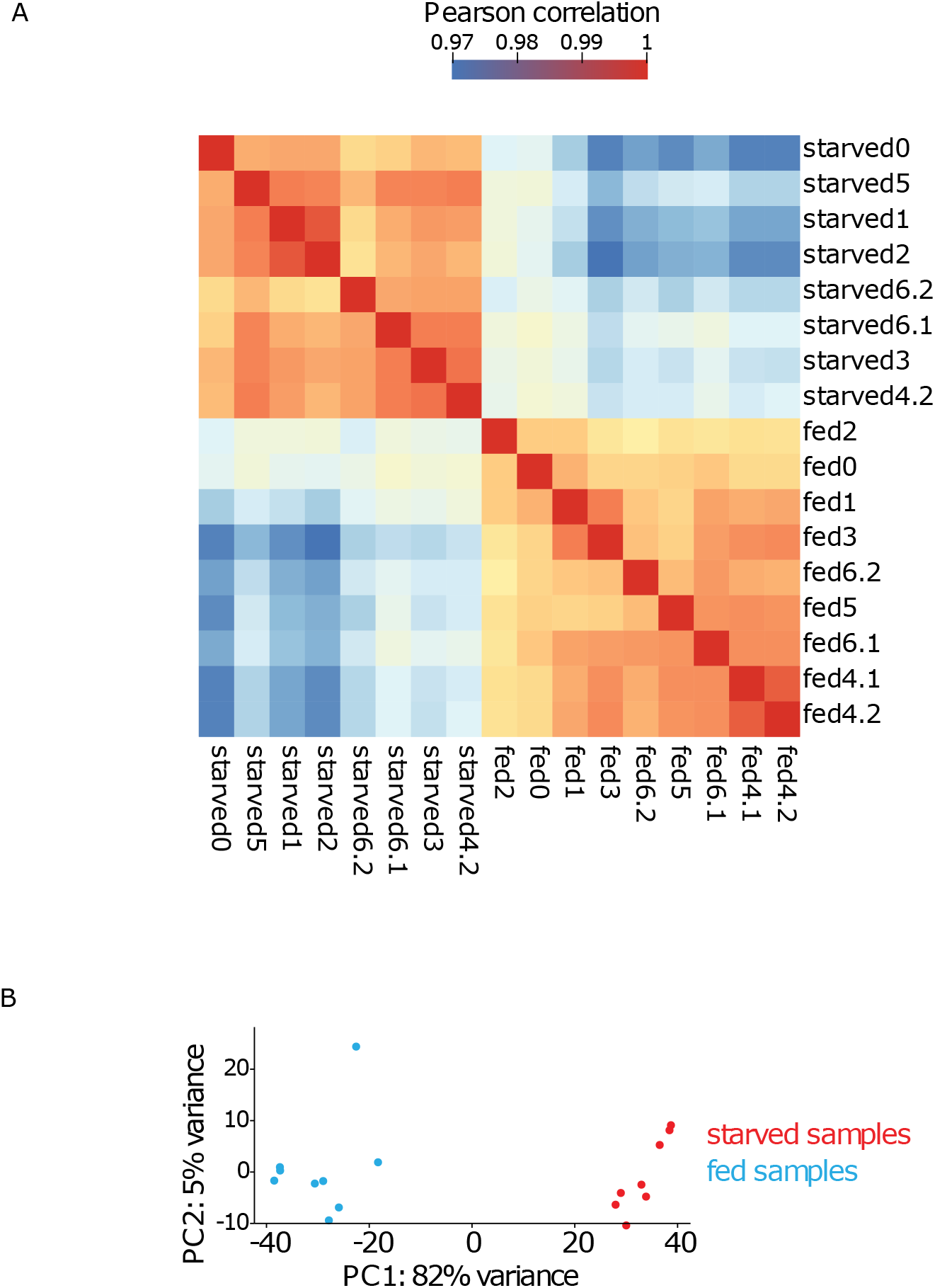
Single-cell RNA-seq reproducibly captured the starvation response. Related to Fig. 1 and 2. (A) Pearson correlation coefficient between each pair of samples (total: 9 fed samples and 8 starved samples) reveals reproducibility among fed and starved replicates as well as differences between fed and starved conditions. (B) Principal component analysis (PCA) of starved and fed pseudobulk samples. The first principal component is correlated with fed vs. starved and explains 82% of the variance, indicating a robust effect of nutrient availability on gene expression.

**Sup. Figure 4.**
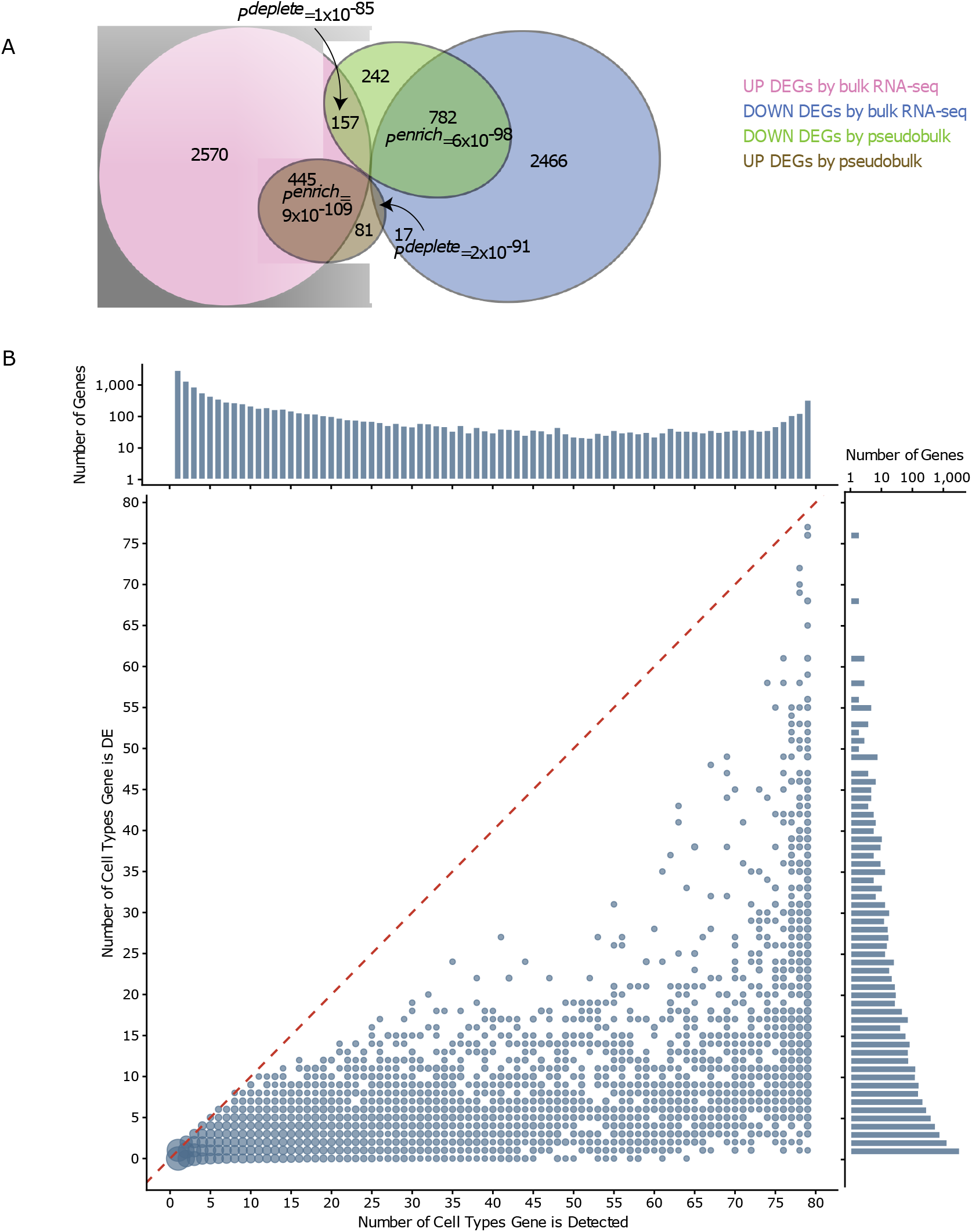
scRNA-seq data are corroborated by published bulk RNA-seq results and reveal that gene expression is detected more broadly than differential expression. Related to Fig. 2. (A) Overlap among genes up and downregulated in starved vs. fed conditions in pseudobulk and a previous bulk mRNA-seq study (Webster *et al*, 2018). Fed and starved samples were collected 18 h and 24 h after bleach, with and without food, respectively, in the bulk mRNA-seq study, which are the same fed and starved time points as in our single-cell RNA-seq study. Hypergeometric tests were done to assess statistical significance of overlaps, with the background set being protein-coding genes detected in both scRNA-seq and bulk mRNA-seq. *P^enrich^* is enrichment p-value from the hypergeometric test. *P^deplete^* is 1 minus *P^enrich^*. (B) Bubble plot relating distributions of the number of cell types genes are detected in and the number of cell types genes are differentially expressed in. A one-dimensional histogram is plotted separately for each axis. Dashed red line indicates y = x.

**Sup. Figure 5.**
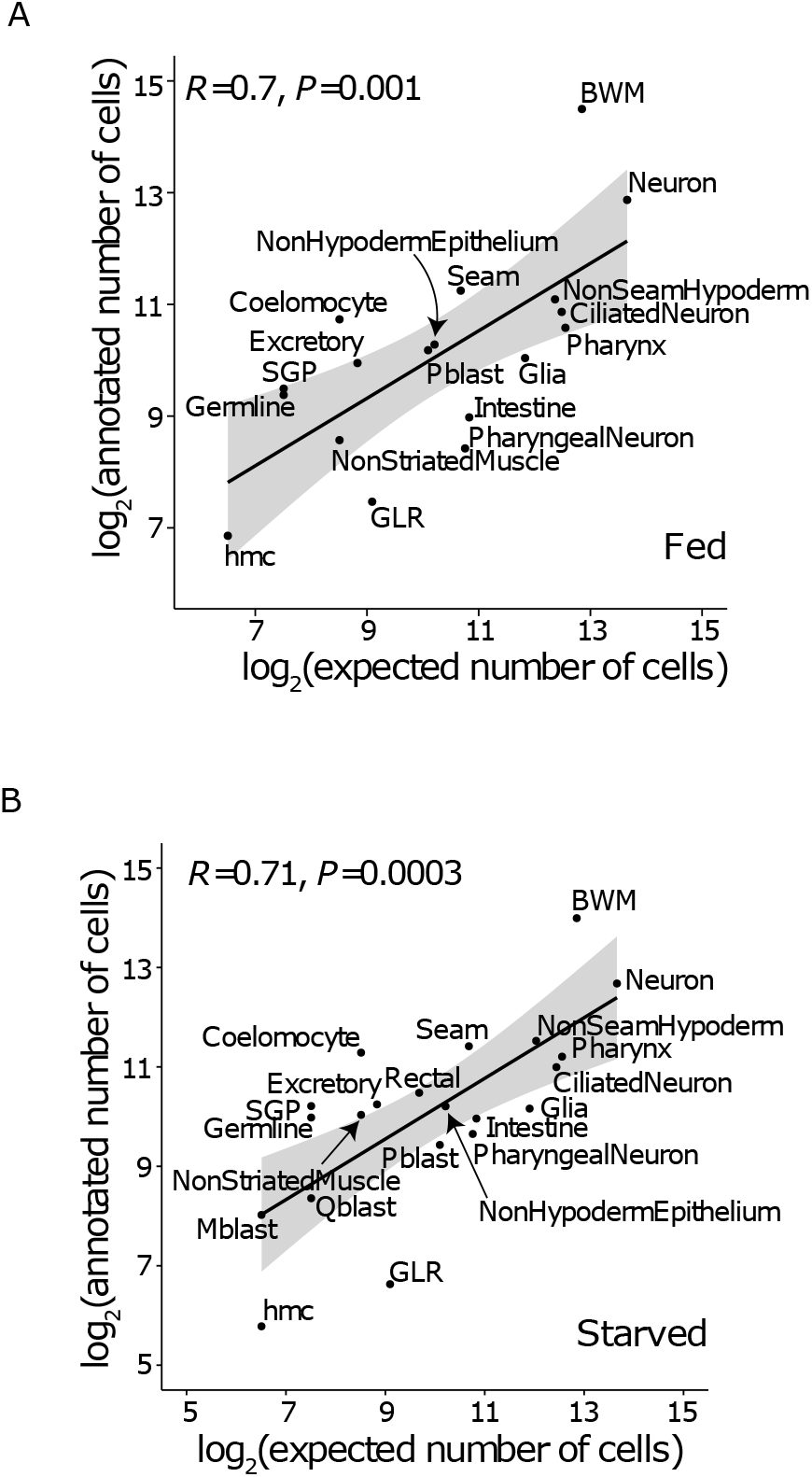
Single-cell RNA-seq reliably identified the major tissues. Related to Figure 3. (A, B) Log_2_ transformed annotated vs. expected number of cells (based on the invariant anatomy of young L1 larvae) for each tissue in fed (A) and starved (B) samples. Pearson correlation coefficients and p-values are indicated. The black line represents a linear regression, and the gray area represents a 95% confidence region. See Sup. Data 2 for a complete list of cell number per tissue in fed and starved conditions.

**Sup. Figure 6.**
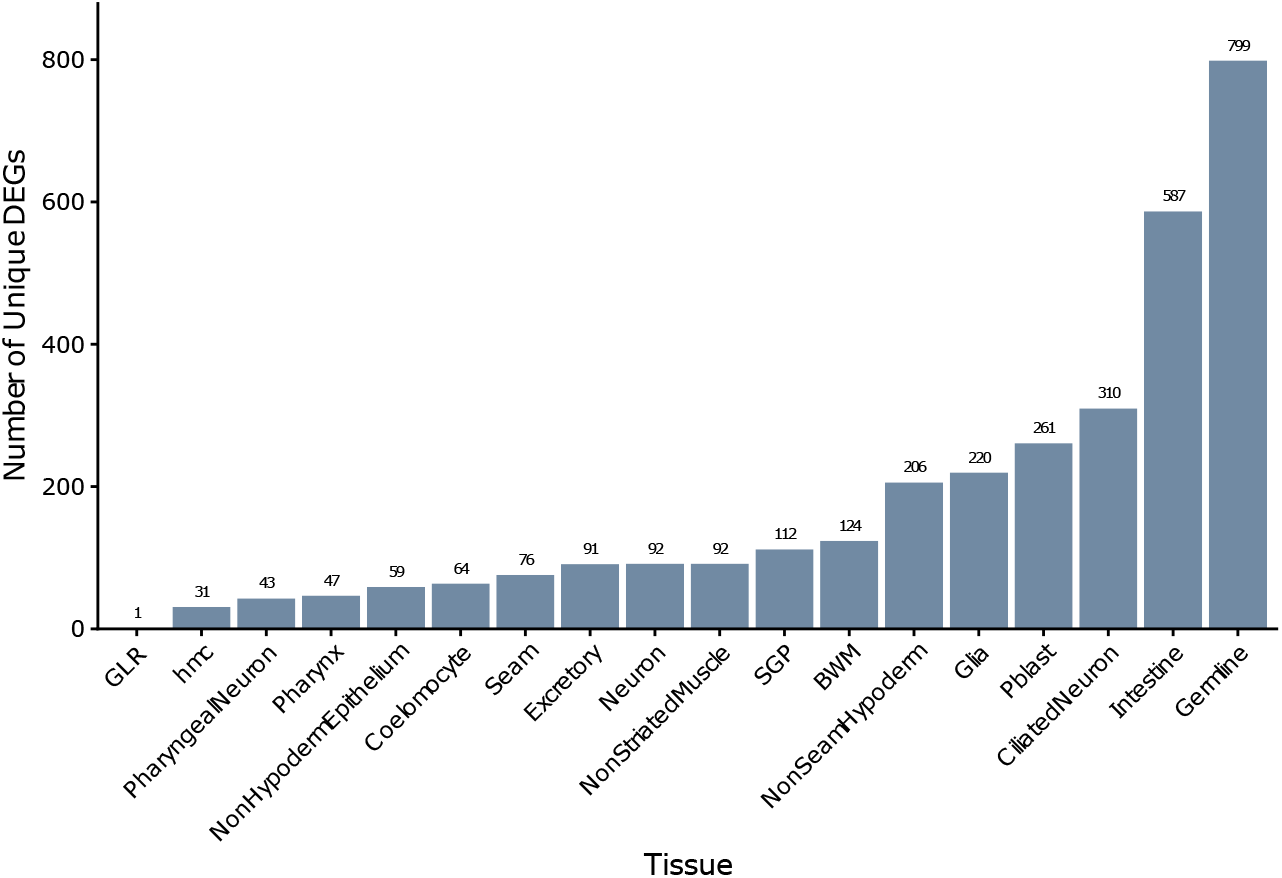
Many genes are differentially expressed in a single tissue. Related to Figure 4. The number of genes that are differentially expressed in a single tissue (‘unique DEGs’) is plotted for each tissue. The germline and intestine are the sites of differential expression for the largest numbers of unique DEGs.

**Sup. Figure 7.**
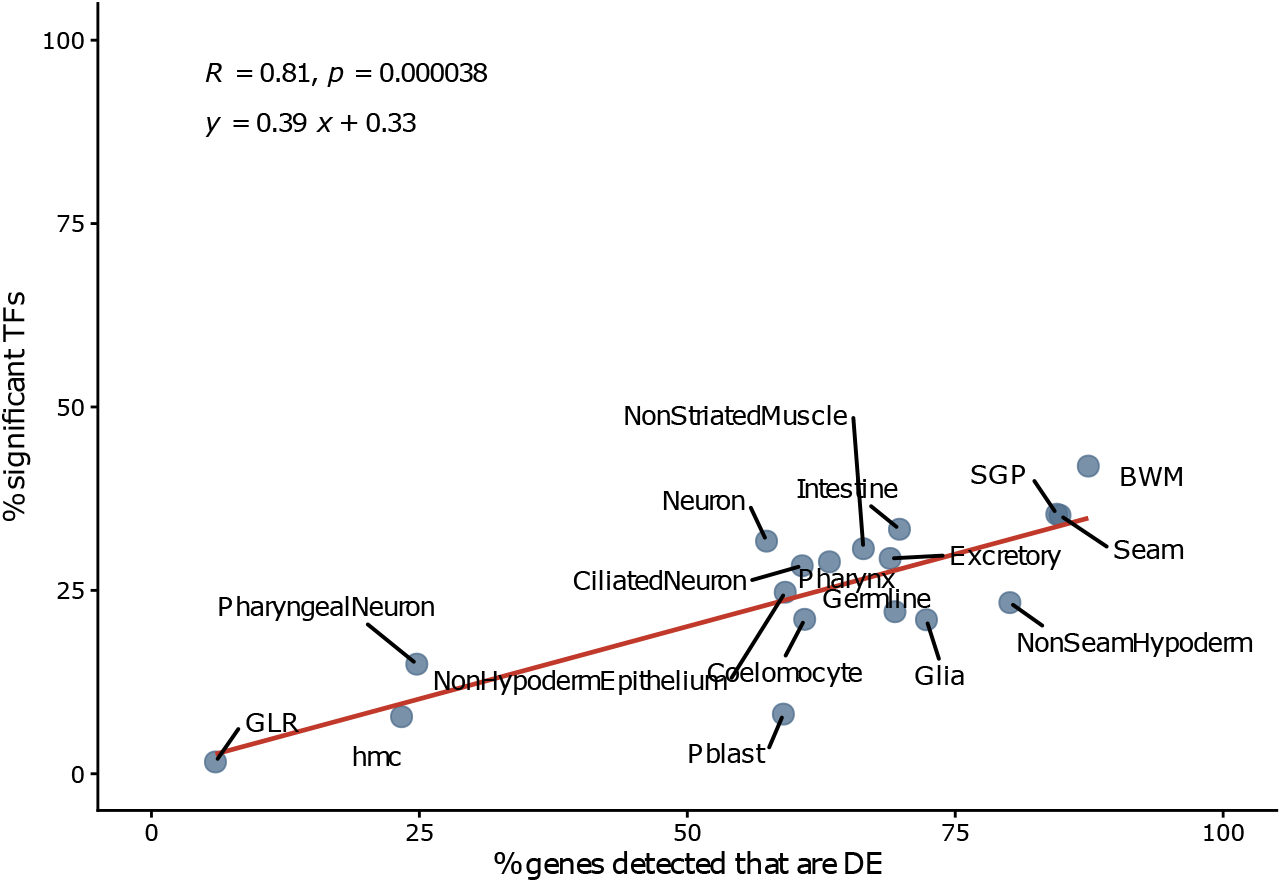
The proportion of TFs with significant tissue-specific differences in activity between fed and starved L1 larvae is proportional to the proportion of genes that differentially expressed in that tissue. The percentage of genes detected in each tissue that are differentially expressed (DE) is plotted against the percentage of transcription factors (TFs) with significant activity estimates per tissue. The red line indicates the linear regression, and the Pearson correlation coefficient, p-value, and slope are given. Related to Figure 6.

**Sup. Figure 8.**
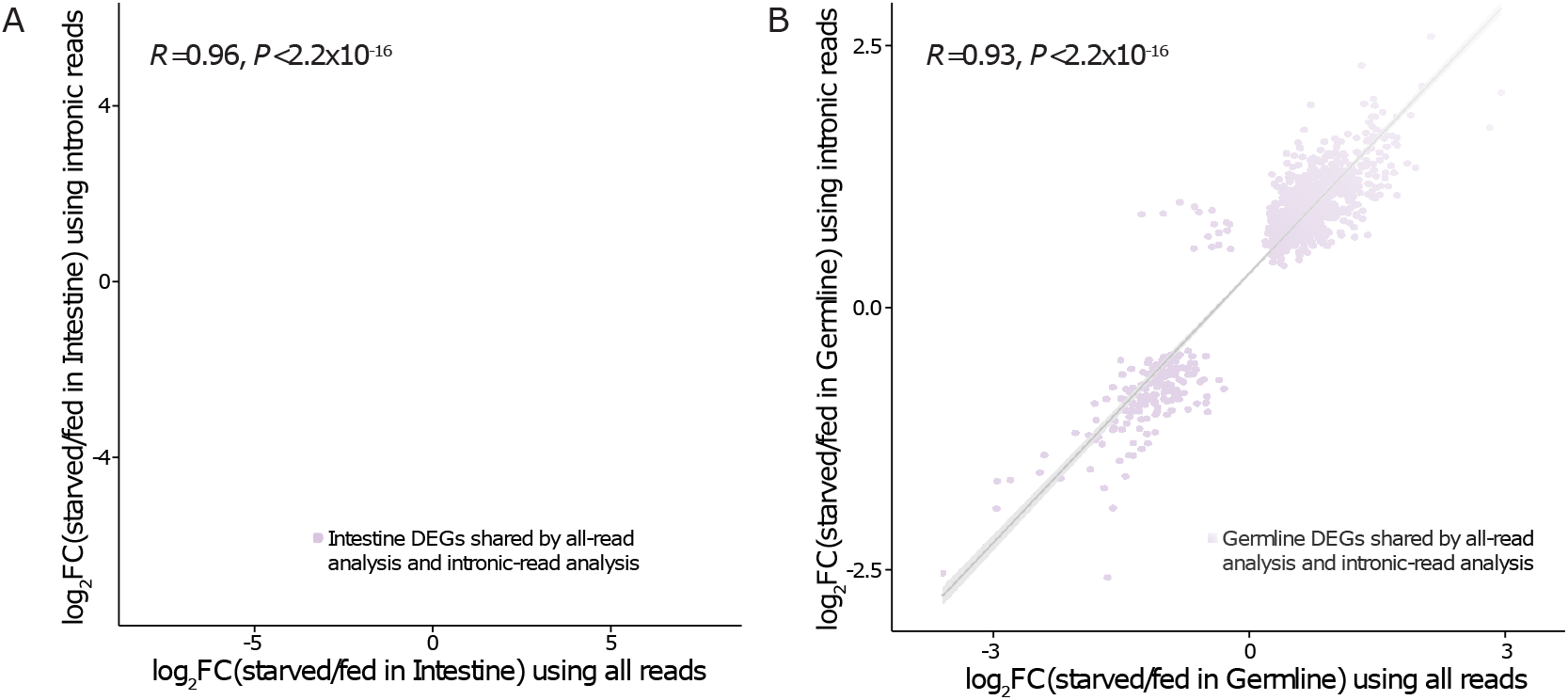
Comparison of differential expression (DE) analysis results using all reads and intronic reads. Related to Figure 7B-E. (A, B) Log_2_fold-change (log_2_FC) of differentially expressed genes (DEGs) using all reads vs. intronic reads for the intestine (A) and germline (B). Pearson correlation coefficient and p-values are indicated. Only DEGs shared by intronic- and all-read analyses were analyzed. Black line represents a linear regression, and the very narrow gray area represents the 95% confidence region. See Sup. Data 4 and 8 for detailed differential expression analysis results using all reads and intronic reads, respectively.

**Sup. Figure 9.**
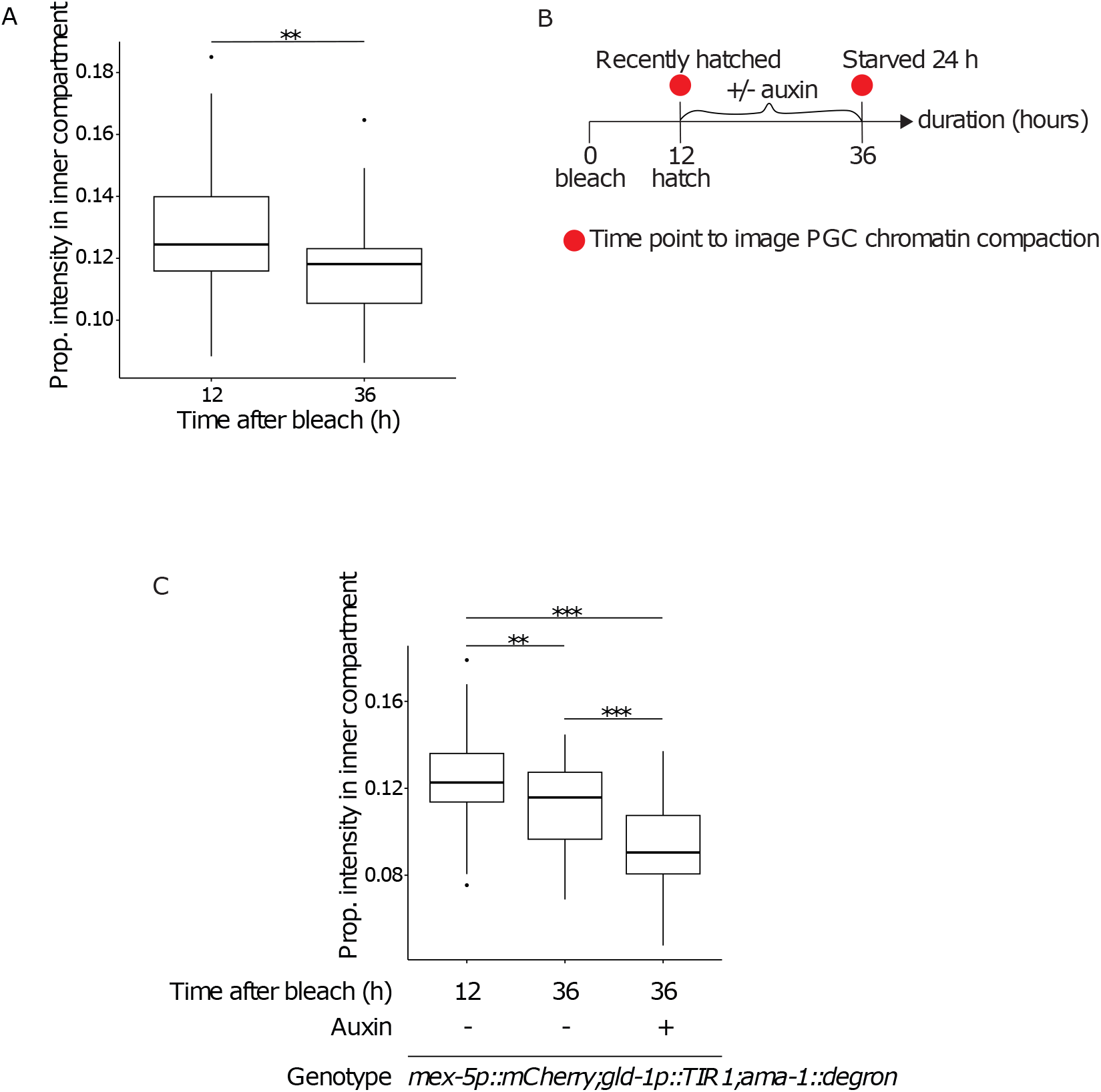
PGC chromatin hyper-compaction due to starvation and severe compaction due to degradation of the large subunit of RNAPII. (A) Proportion of PGC chromatin intensity in the ‘inner compartment’ (11.1% as previously defined (Belew *et al*, 2021)) in recently hatched (12 h after bleach) vs 24 h starved (36 h after bleach) wild-type worms. Number of PGCs: 51 and 67 for recently hatched and 24 h starved worms, respectively. Because Bartlett’s test rejected equal variance between the two conditions, an unpaired t-test was performed without pooling variances. (B) Schematic for the auxin-induced degradation (AID) experiment to degrade AMA-1/RNAPII and image PGC chromatin compaction. (C) Proportion of PGC chromatin intensity in the ‘inner compartment’ (11.1%) in recently hatched (12 h after bleach) vs 24 h starved (36 h after bleach) worms with or without auxin to degrade AMA-1/RNAPII. Number of PGCs: 57 for recently hatched worms, 40 for 24 h starved worms without auxin, and 37 for 24 h starved worms with auxin. Barlett’s test did not reject equal variance among the three groups, so unpaired t-tests were performed with pooled variances. (A, C) **P<0.01; ***P<0.001.

**Sup. Table 1.**
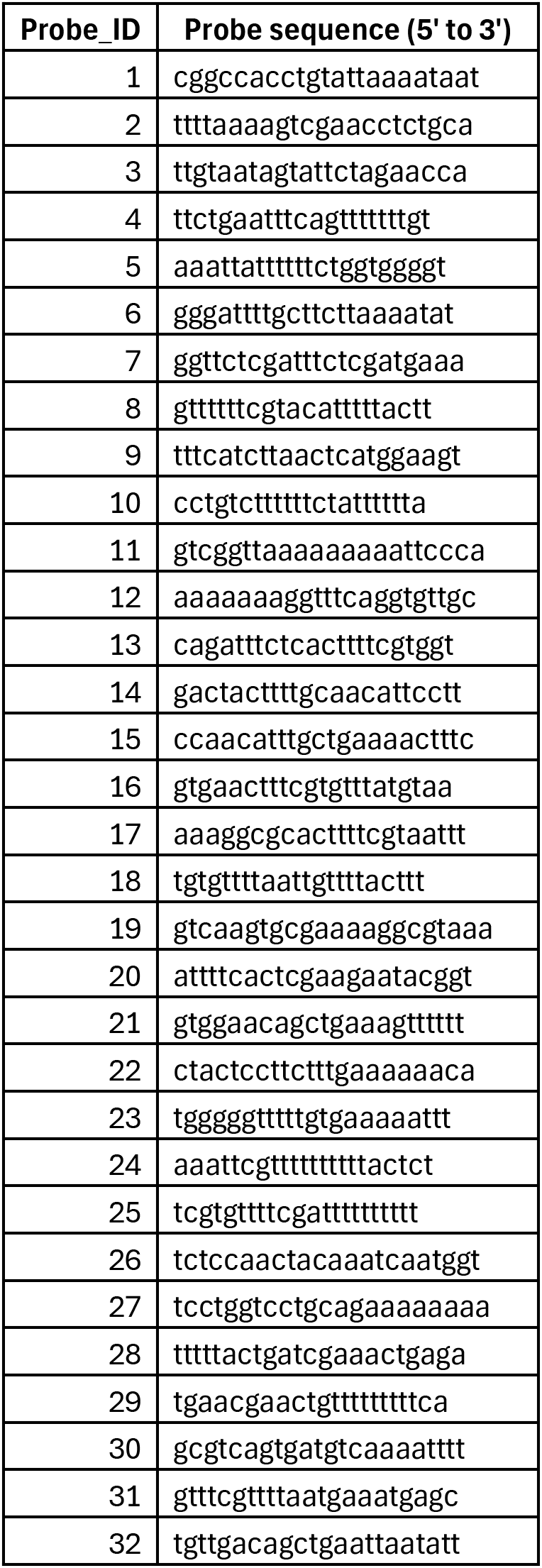
5’ to 3’ sequence of 32 *aak-1/AMPK* intronic probes for smFISH. Related to Figure 6A-D.

**Sup. Data 1.** Post-quality-control sample metrics. UMI: Unique Molecular Identifier. Proportion (Prop.) genome was calculated by dividing number (No.) detected genes by the total number of genes in WS286 (46,926). Prop. protein-coding genome was calculated by dividing No. detected protein-coding genes by the total number of protein-coding genes in WS286 (19,983). Nine fed samples and eight starved samples passed quality control. Metrics for fed, starved, and total were calculated by aggregating all fed, all starved, and all samples, respectively. Related to Fig. 1.

**Sup. Data 2.** Information about marker genes used to annotate each cell type, how cell types were aggregated into tissues, cell type and tissue abbreviation rules, and the expected number of cells in each tissue in fed and starved conditions. Related to Figs. 1, 3, and S5.

**Sup. Data 3.** Cell-type-level differential expression analysis results. Transcript abundance and fold-changes for each gene in each cell type can also be plotted online (https://jingxianchen.shinyapps.io/l1_sc/). Related to Fig. 2.

**Sup. Data 4.** Tissue-level differential expression analysis results. Transcript abundance and fold-changes for each gene in each tissue can also be plotted online (https://jingxianchen.shinyapps.io/l1_sc/). Related to Fig. 4.

**Sup. Data 5.** Gene Ontology (GO) term analysis results. Related to Fig. 6.

**Sup. Data 6.** Results from estimation of transcription factor activities using *Cel*EsT.

**Sup. Data 7.** KEGG pathway enrichment analysis results for germline.

**Sup. Data 8.** Differential expression analysis results using intronic reads for intestine and germline.

